# A redox-regulated, heterodimeric NADH:cinnamate reductase in *Vibrio ruber*

**DOI:** 10.1101/2023.03.04.531086

**Authors:** Yulia V. Bertsova, Marina V. Serebryakova, Victor A. Anashkin, Alexander A. Baykov, Alexander V. Bogachev

**Affiliations:** Belozersky Institute of Physico-Chemical Biology, Lomonosov Moscow State University, Moscow 119234, Russia

**Keywords:** anaerobic respiration, cinnamate, enzyme regulation, *Vibrio*

## Abstract

Genes of putative reductases of α,β-unsaturated carboxylic acids are abundant among anaerobic and facultatively anaerobic microorganisms, yet substrate specificity has been experimentally verified for few encoded proteins. Here, we co-produced in *Escherichia coli* one such protein of the marine bacterium *Vibrio ruber* (GenBank SJN56021; annotated as urocanate reductase) with *Vibrio cholerae* flavin transferase. The isolated protein (named Crd) is a heterodimer of the SJN56021-encoded subunit CrdB (*NADH:flavin, FAD binding 2*, and *FMN bind* domains) and an additional subunit CrdA (SJN56019, a single *NADH:flavin* domain) that interact via their *NADH:flavin* domains (Alphafold2 prediction). Each domain contains a flavin group (three FMNs and one FAD in total), one of the FMN groups being linked covalently by the flavin transferase. Crd readily reduces cinnamate, *p*-coumarate, caffeate, and ferulate under anaerobic conditions with NADH or methyl viologen as the electron donor, is moderately active against acrylate and practically inactive against urocanate. The reduction reactions started by NADH demonstrated a time lag of several minutes, suggesting a redox regulation of Crd activity. The oxidized enzyme is inactive, which apparently prevents production of reactive oxygen species under aerobic conditions. Our findings identify Crd as a regulated NADH-dependent cinnamate reductase, apparently protecting *V. ruber* from cinnamate poisoning.

**Abbreviated Summary:** The genome of the marine bacterium *Vibrio ruber* encodes a heterodimeric NADH-dependent cinnamate reductase, apparently protecting *V. ruber* from poisoning by cinnamate and its derivatives. The reductase contains four flavin groups, one being linked covalently, and appears to be redox-regulated. Oxidized enzyme is inactive, which apparently prevents production of reactive oxygen species under aerobic conditions.

## 1 INTRODUCTION

NADH:2-enoate reductases (EC 1.3.1.31) capable of reducing α,β-unsaturated carboxylic acids, such as fumaric, cinnamic, and acrylic acids, are abundant among anaerobic and facultatively anaerobic microorganisms (Bertsova et al., 2020; 2022; Besteiro et al., 2002; Tischer et al., 1979). All known 2-enoate reductases are formed by single polypeptides but are divided into two non-homologous groups according to their variable domain composition. The three-domain reductases invariably contain a *FAD binding 2* domain (Pfam ID: PF00890), which has a noncovalently bound FAD prosthetic group and is the place of carbonic acid reduction. NADH is oxidized in *OYE*-*like* (PF00724) or *FAD binding 6* (PF00970) domain having noncovalently bound FMN or FAD, respectively (Bertsova et al., 2020; Besteiro et al., 2002). The third domain (commonly *FMN bind*; PF04205) mediates electron transfer between the above domains via a flavin group (FMN) which is covalently bound by a phosphoester bond (Bertsova et al., 2014; 2022; Serebryakova et al., 2018).

Many bacteria of the class Clostridia have a two-domain 2-enoate reductase (Rohdich et al., 2001; Tischer et al., 1979), formed by *OYE*-*like* and *Pyr redox 2* (PF07992) domains. The reductases of this group contain a [4Fe-4S] cluster and noncovalently bound FAD and FMN in a 1:1:1 ratio as prosthetic groups (Caldeira et al., 1996; Kuno et al., 1985). In saccharolytic clostridia, two-domain 2-enoate reductases convert a broad range of 2-enoates, whereas the enzymes from proteolytic clostridia are highly specific and convert only cinnamate and its derivatives (Giesel & Simon, 1983; Mordaka et al., 2018). All known 2-enoate reductases do not appear to be regulated at activity level, except by substrates and products.

The genome of the red facultatively anaerobic marine bacterium *Vibrio ruber* (Shieh et al., 2003) encodes a polypeptide formed by 806 amino acid residues and annotated as urocanate reductase in GenBank (SJN56021) or urocanate reductase precursor in UniProt (A0A1R4LHH9). This polypeptide, which we designate as CrdB, based on the substrate specificity of its containing enzyme (see below), encompasses the abovementioned *FAD binding 2* and *FMN bind* domains (Fig. 1A). Five putative substrate-binding residues, identified in the *FAD binding 2* domains of CrdB and acrylate reductase of *Vibrio harveyi* by comparison with fumarate reductase (Bertsova et al., 2022; Bogachev et al., 2012; Light et al., 2019) (Fig. 1B), are identical or very similar, suggesting that CrdB is an acrylate reductase. The *FMN bind* domain of CrdB, like its counterparts in known 2-enoate reductases, contains the motif DALSGAS_257_ recognized by flavin transferase for covalent FMN attachment to the serine residue (Bogachev et al., 2018). Noteworthy, the gene *apb*E encoding the flavin transferase (GenBank ID: SJN56016) is adjacent to *crd*B in one of the two *V. ruber* chromosomes (Fig. 1C). Altogether, these characteristics of Crd classify it as a three-domain 2-enoate reductase with a likely acrylate reductase activity.

**Figure 1.**
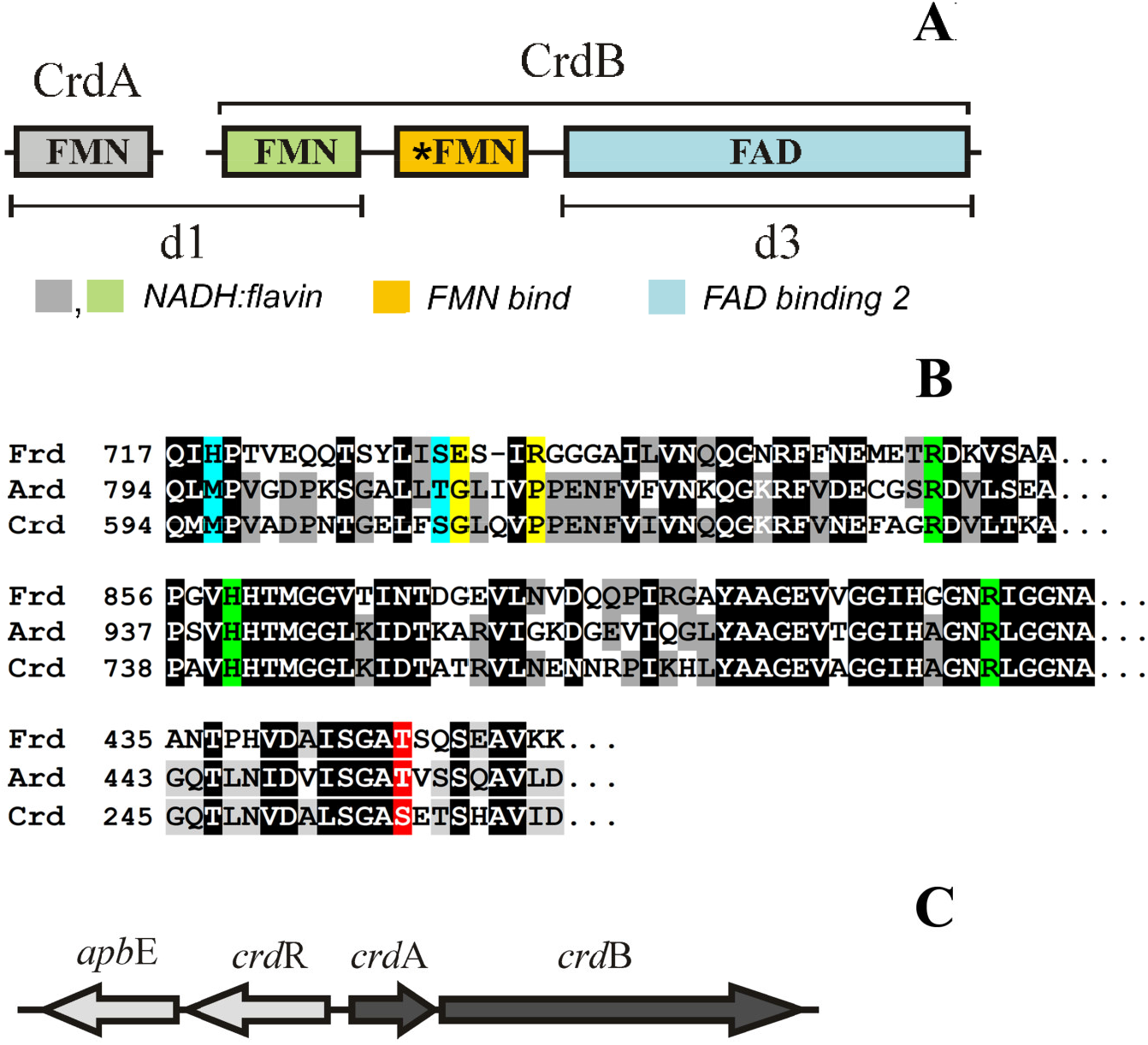
Bioinformatic description of *V. ruber* Crd and its genes. (A) The domain composition of the polypeptides CrdA and CrdB forming the Crd heterodimer. The *NADH:flavin, FMN bind*, and *FAD binding 2* domains (residues 1–180, 190–275, and 280–800, respectively) are indicated by different colors. Putative redox-active prosthetic groups are shown inside the boxes. The asterisk denotes the covalently bound flavin. (B) Sequence alignment of the fumarate reductase from *K. pneumoniae* (Frd, UniProt accession number B5XRB0), acrylate reductase from *V. harveyi* (Ard, P0DW92), and CrdB protein from *V. ruber* (GenBank accession number SJN56021) with Clustal (Sievers & Higgins, 2018). The three parts of the alignment shown contain the amino acid residues involved in fumarate C1- and C4-carboxylate binding in Frd (marked in blue and green, respectively), the proton transfer to fumarate (marked in yellow) in Frd, and covalent bonding with FMN in all shown proteins (marked in red). (C) The arrangement of the *crd*-associated genes in a *V. ruber* chromosome. ApbE, putative FAD:protein FMN transferase; CrdR, putative transcription regulator.

However, the CrdB protein differs from established three-domain 2-enoate reductases by its third, *NADH:flavin* domain (PF03358) replacing the NADH-oxidizing domains *OYE*-*like* or *FAD binding 6* (Fig. 1A). Furthermore, the *crd*-operon of *V. ruber* contains, upstream of the *crd*B gene, the gene *crd*A encoding a protein (SJN56019) (Fig. 1C) formed by a single *NADH:flavin* domain that is homologous (51 % identity, 65 % similarity) to the similar domain of CrdB. As *NADH:flavin* domains can dimerize (Koike et al., 1998), this observation raised an intriguing possibility of CrdB functioning as heterodimer.

Keeping in mind the difficulty in the theoretical prediction of the enzymatic activities and substrate specificities of NADH:2-enoate reductases, we have produced the *V. ruber* Crd protein in *Escherichia coli* cells and characterized the isolated protein. The data reported below identify Crd as a regulated NADH-dependent reductase active against cinnamic acid and its various derivatives.

## 2 RESULTS

### 2.1 Production and characterization of *V. ruber* Crd

The CrdAB operon of *V. ruber* genomic DNA was amplified and cloned into an expression vector that added a 6×His tag at the C-terminus of CrdB. The *FMN bind* domain of CrdB harbors the sequence DALSGA**S**_**257**_, similar to the flavinylation motif Dxx(s/t)gA(**T/S**) recognized by flavin transferase for covalent attachment of FMN via a threonine or serine hydroxyl (Bogachev et al., 2018). To test whether the predicted flavinylation really occurs, the *crd*-operon genes were expressed in *E. coli* cells in the presence or absence of the auxiliary pΔhis3 plasmid (Bertsova et al., 2013) that encodes the flavin transferase ApbE capable of flavinylating *FMN bind* domains in various proteins (Bogachev et al., 2018). The recombinant fCrd and aCrd proteins produced in the ApbE-containing and ApbE-lacking cells, respectively, were isolated by metal affinity chromatography. The final protein preparations were of intense yellow color, and their visible spectra (Fig. 2A) were characteristic of flavoproteins.

**Figure 2.**
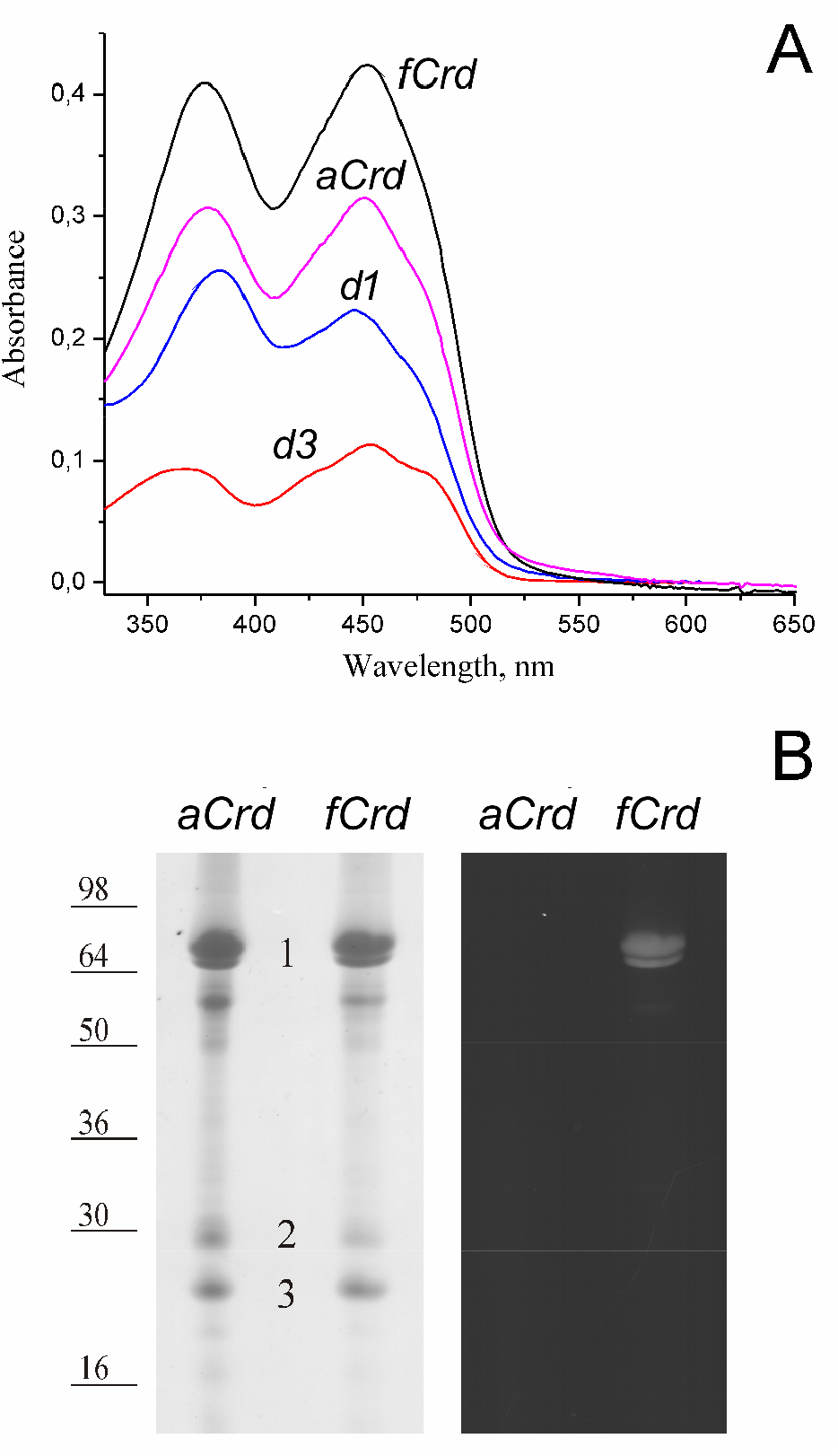
Characterization of the Crd preparations. (A) Electronic absorption spectra of the full-length and truncated Crd preparations at 10 μM concentrations. Full-length Crd was produced in ApbE-containing or ApbE-lacking *V. ruber* cells (fCrd and aCrd, respectively). (B) SDS-PAGE of fCrd and aCrd. The gel was stained with Coomassie Blue (left image) or scanned under illumination at 473 nm without staining (right image). The protein load was 5 μg per lane. Bars with numbers on the left side denote the positions and molecular masses of marker proteins. The protein bands identified by MALDI-MS analysis are marked by numbers.

SDS-PAGE of both Crd preparations revealed one major and three–four minor protein bands (Fig. 2B, left image). Bands 1 and 3 were identified by the MALDI-MS analysis after trypsin digestion as *V. ruber* CrdB (sequence coverage of 79%) and CrdA (sequence coverage of 95%), respectively. Band 2 belonged to *E. coli* peptidyl-prolyl *cis*-*trans* isomerase SlyD, which exhibits high intrinsic affinity to Ni-agarose and is a common contamination of the 6×His-tagged proteins isolated from *E. coli* (Bolanos-Garcia & Davies, 2006). Band 1 derived from fCrd fluoresced when illuminated at 473 nm, indicating covalently bound flavin (Fig. 2B, right image). No bound flavin was detected by this method in aCrd. These findings provide convincing evidence for ApbE-mediated covalent flavinylation of CrdB. It should be noted that the non-covalently bound flavins, evidently present in both Crd preparations (Fig. 2A) are removed from the proteins during SDS-PAGE.

The CrdB flavinylation site was identified by MALDI-MS and MS/MS after trypsinolysis. The peptide I_242_IEGQTLNV<u>DALSGA</u>**<u>S</u>**ETSHAVIDGVAK_269_ containing the underlined flavinylation motif should have demonstrated the monoisotopic MH^+^ masses of 2795.3 and 3233.5 in the non-flavinylated and flavinylated forms, respectively. Both predicted signals were indeed observed on the mass-spectrum of the tryptic digest of fCrdB. The presence of the former signal may reflect either incomplete modification or partial loss of the flavin group during SDS-PAGE and trypsin proteolysis. No other candidates for flavinylated peptides were observed in the tryptic digest of CrdB and CrdA by a peptide fingerprint Mascot search.

The MS/MS analysis of the peptide with *m/z* of 3233.5 (Fig. 3) indicated that the main signals in the spectrum of fragmentation (*m/z* of 2794 and 2776) resulted from FMN loss in the full or dehydrated form, respectively, depending on which of the two covalent bonds, C–O or O– P, connecting the FMN moiety to the serine residue is broken. Greater intensity of the signal with *m/z* of 2776 indicated that the former bond is broken with greater propensity, resulting in serine dehydration (Bertsova et al., 2021). Less intense but still reliable peaks in the spectrum starting from the fragment with *m/z* = 2776 also matched a series of the *y*-ions generated by double breaks during the fragmentation (Fig. 3, red letters). Here, mass difference for the Ser257 position (69 Da) corresponded to the dehydrated serine, demonstrating that fCrdB produced in ApbE-containing cells harbors an FMN residue covalently bonded to Ser257 of the predicted flavinylation motif in the *FMN bind* domain.

**Figure 3.**
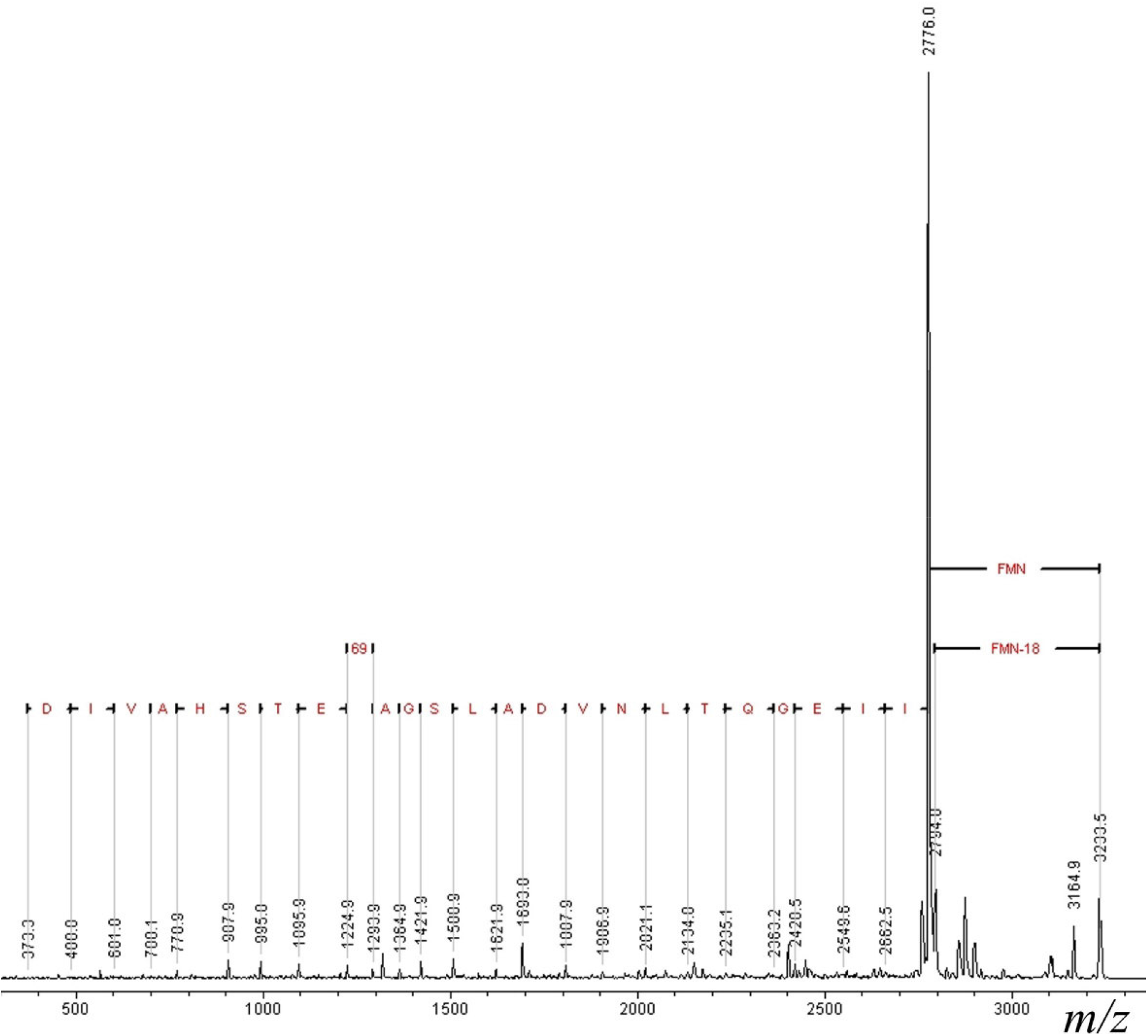
The MS/MS spectrum of the fCrdB peptide with *m/z* of 3233.5. The protein sequence deduced from the series of *y*-ions is shown by red letters.

The non-covalently bound flavins were extracted from Crd by TFA and separated by HPLC (Fig. 4). Both the fCrd and aCrd preparations were found to contain non-covalently bound FAD and FMN at a ratio of approximately 1:2.

**Figure 4.**
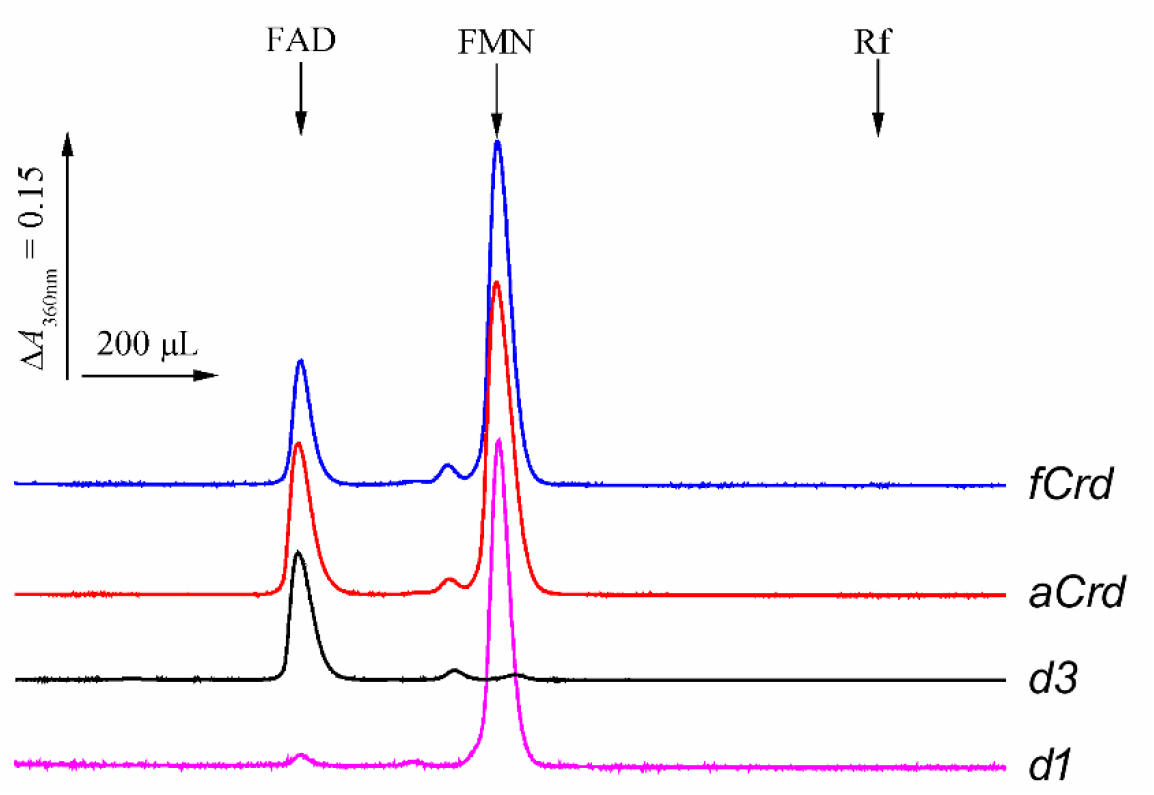
HPLC separation of non-covalently bound flavins in Crd and its deletion variants. The retention volumes for authentic FAD, FMN, and riboflavin (Rf) are indicated by arrows.

To determine their domain localization, we prepared the genetic constructs encoding separately two *NADH:flavin* domains of CrdA and CrdB (d1 in Fig. 1A) or the *FAD binding 2* domain of CrdB (d3 in Fig. 1A). The isolated corresponding proteins exhibited the spectra characteristic of flavoproteins (Fig. 2A). The flavins present in them were again extracted and identified by HPLC (Fig. 4). The non-covalently bound FMN was exclusively localized in the d1 fragment (*NADH:flavin* domains) whereas FAD was found in the d3 fragment (*FAD binding 2* domain). The evident corollary is that CrdAB contains two non-covalently bound FMN molecules in the oxidoreductase *NADH:flavin* domains, one non-covalently bound FAD molecule in the *FAD binding 2* domain, and one covalently bound FMN residue in the *FMN bind* domain.

### 2.2 The specificity of Crd for naturally occurring α,β-unsaturated carbonic acids

The similarity of the putative substrate-binding residues in the *FAD binding 2* domains of Crd and the acrylate reductase Ard of *V. harveyi* (Fig. 1B) raised the possibility that Crd also possesses acrylate reductase activity. Indeed, the Crd preparation obtained by *crd*AB-*apb*E coexpression (fCrd) could reduce acrylate using reduced MV as the electron donor, but at a much lower rate (1.3 s^-1^) compared with that of a specific acrylate reductase (19 s^-1^ (Bertsova et al., 2022)). Conversely, the *K*_m_ value was much greater for fCrd (770 versus 16 μM), making its catalytic efficiency *k*_cat_/*K*_m_ 640 times lower (0.0019 μM^-1^ s^-1^) compared with the acrylate reductase (1.2 μM^-1^ s^-1^ (Bertsova et al., 2022)). fCrd exhibited low but measurable activity against fumarate but was virtually inactive with methacrylate, crotonate, and urocanate (Table 1).

**Table 1.**
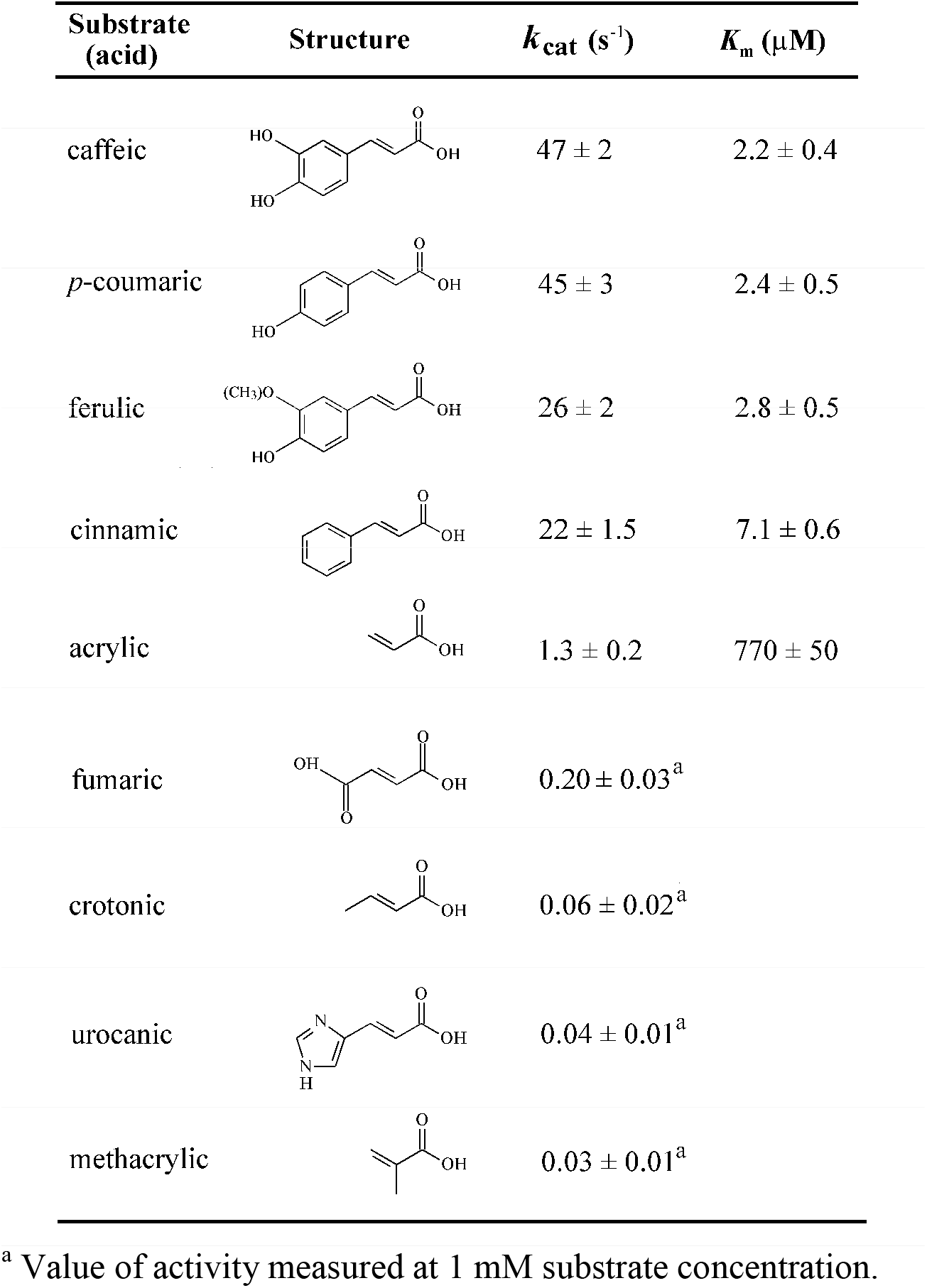
The reductase activity of fCrd against a series of α,β-unsaturated carbonic acids measured under anaerobic conditions with 100 μM reduced MV as the electron donor.

| Substrate (acid) | Structure | $k_{\text{cat}} (\text{s}^{-1})$ | $K_m (\mu\text{M})$ |
| --- | --- | --- | --- |
| caffeic | | $47 \pm 2$ | $2.2 \pm 0.4$ |
| *p*-coumaric | | $45 \pm 3$ | $2.4 \pm 0.5$ |
| ferulic | | $26 \pm 2$ | $2.8 \pm 0.5$ |
| cinnamic | | $22 \pm 1.5$ | $7.1 \pm 0.6$ |
| acrylic | | $1.3 \pm 0.2$ | $770 \pm 50$ |
| fumaric | | $0.20 \pm 0.03^a$ | |
| crotonic | | $0.06 \pm 0.02^a$ | |
| urocanic | | $0.04 \pm 0.01^a$ | |
| methacrylic | | $0.03 \pm 0.01^a$ | |

<sup>a</sup> Value of activity measured at 1 mM substrate concentration.

In contrast, cinnamate and its hydroxy derivatives were readily reduced by fCrd (*k*_cat_ of 22–47 s^-1^) and demonstrated low *K*_m_ values of 2.2–7.1 μM (Table 1). These findings classify fCrd as a cinnamate reductase that is also active with other natural α,β-unsaturated cinnamate derivatives.

To identify the product of cinnamate reduction by fCrd with MV as the electron donor, we extracted the organic material from the reaction mixture with diethyl ether before and after complete conversion of 100 μM cinnamate and separated the extracts by HPLC. Complete conversion of cinnamate to phenylpropionate was observed (Fig. 5A,B). The identification of the carbonic acids was supported by the observation that the peak heights on the elution profiles monitored at 210 and 258 nm were nearly equal for cinnamate but differed approximately 40-fold for phenylpropionate, in accordance with the UV-spectra of these compounds (Fig. 5C). These findings indicate that Crd reduces the α,β double bond in (hydroxy)cinnamic acids to yield their saturated derivatives. fCrd activity in the reverse reaction of phenylpropionate oxidation with phenazine methasulfate and dichlorophenolindophenol as electron acceptors was measurable but quite low (0.1 s^-1^).

**Fig. 5.**
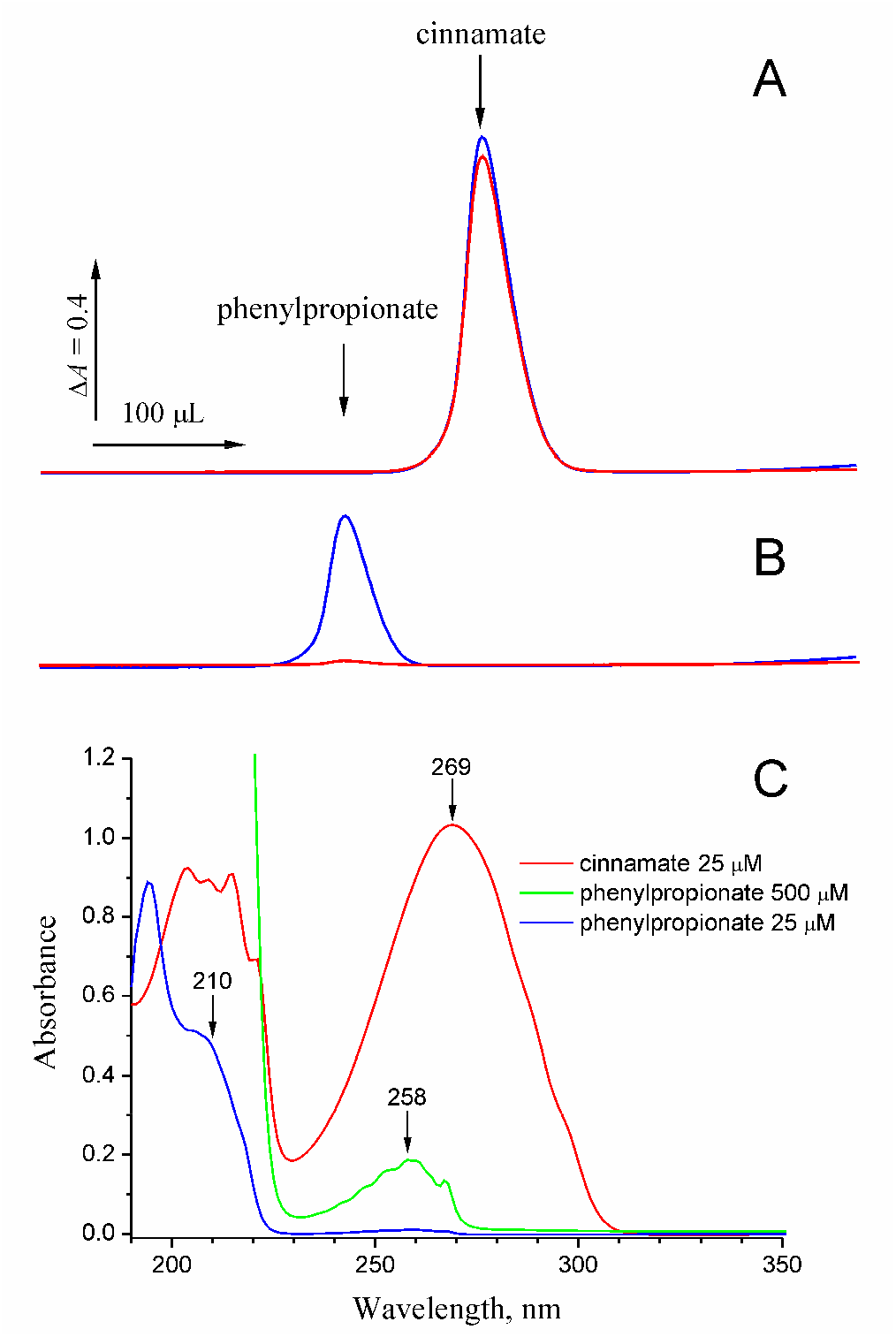
HPLC identification of the product of cinnamate reduction by fCrd. Cinnamate (0.1 mM) was reacted with 10 mM sodium dithionite and 50 μM MV in the presence of 0.5 μM fCrd. (A) The reaction mixture before fCrd addition. (B) Same after a 30-min incubation with fCrd. The elution was monitored at 210 (blue curve) and 258 nm (red curve). The elution volumes for authentic cinnamate and phenylpropionate are indicated by arrows. (C) Electronic absorption spectra of cinnamate and phenylpropionate in 10 mM potassium phosphate buffer (pH 6.5).

### 2.3 Time-dependent activation of Crd by NADH

The presence of the *NADH:flavin* domains in CdrAB (Fig. 1A) suggested that NADH is the natural donor of the redox equivalents for this enzyme. As ferulate, caffeate, and *p*-coumarate absorb light at 340 nm, the NADH oxidase activity was tested with non-absorbing cinnamate as the electron acceptor. As the red trace in Fig. 6A highlights, the reaction initiated by cinnamate addition under anaerobic conditions demonstrated a linear NADH oxidation curve at a relatively high rate of 15 s^-1^. However, changing the order of the additions affected the reaction time course dramatically. For the reaction started by fCrd, the progress curve demonstrated a profound lag in the NADH oxidation (blue trace in Fig. 6A). Although the initial slope was quite low, the final slope was similar to that for the red trace described above and likely corresponded to fully activated Crd. A similar time lag was observed in the reaction started by NADH.

**Figure 6.**
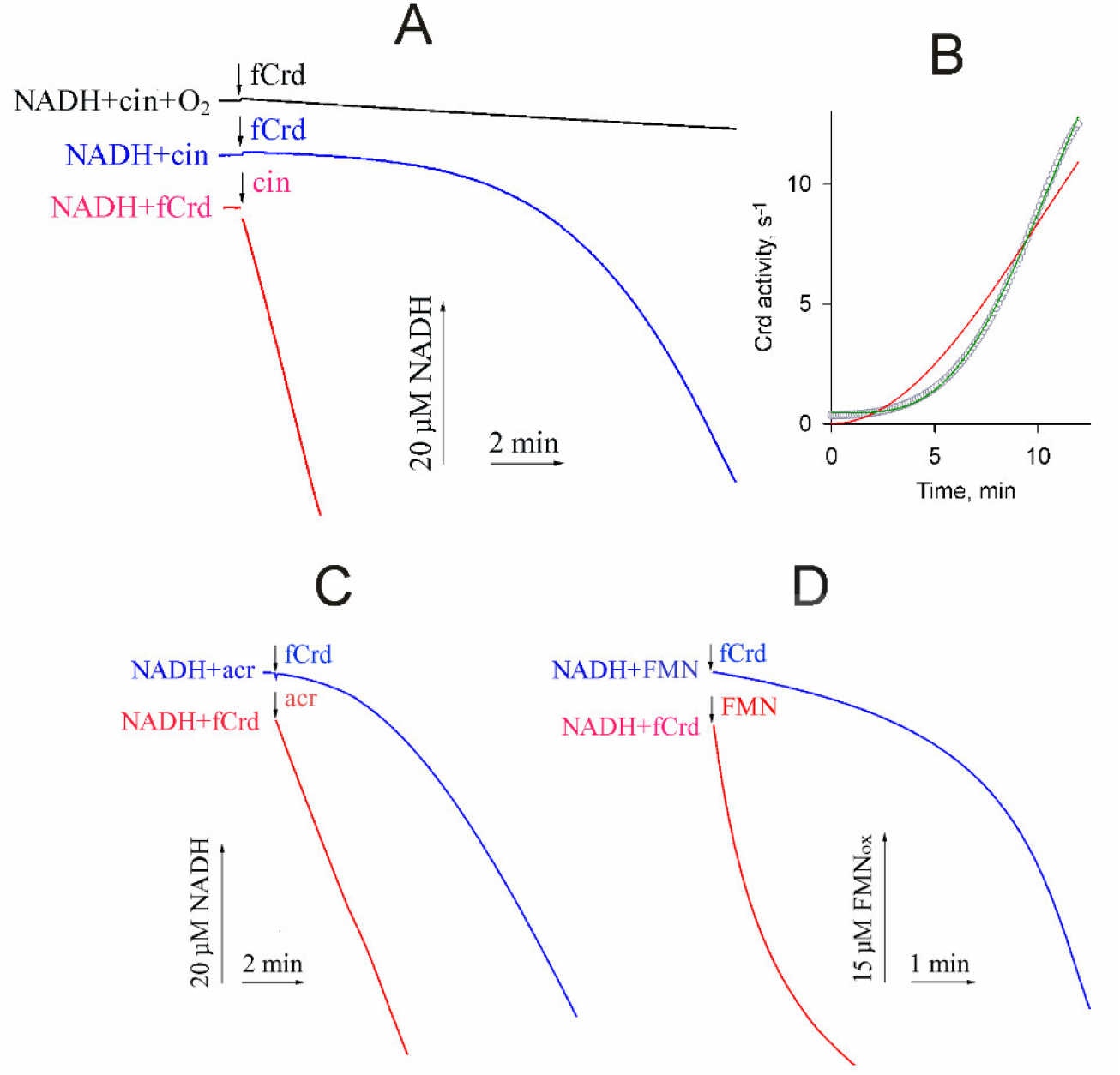
The typical traces of fCrd-catalyzed NADH-dependent reactions. (A) Cinnamate-supported NADH oxidation. The final reaction mixture contained 120 μM NADH, 50 μM cinnamate, and 30 nM fCrd. The reaction was carried out under anaerobic conditions (blue and red traces) and started by either NADH (red trace) or fCrd (blue trace). The black trace was obtained under aerobic conditions for the reaction started by fCrd. (B) The time dependence of Crd activity derived from the time-course of cinnamate reduction in the reaction started by Crd addition (blue curve on panel A). Small circles show the activity values determined at 100 time points as the slope of the tangent (-*d*[NADH]/*d*t) using Matlab (The Mathworks, Inc.). The red and green lines show the best fits for two- and three-step kinetic models, respectively. (C) Anaerobic acrylate-supported NADH oxidation. The final reaction medium contained 120 μM NADH, 1 mM acrylate, and 150 nM Crd. The reaction was started by acrylate (red trace) or NADH (blue trace). (D) Anaerobic NADH-supported FMN reduction. The final reaction medium contained 120 μM NADH and 50 μM FMN. The fCrd concentration was 15 nM (red trace, reaction started by FMN) or 60 nM (blue trace, reaction started by fCrd).

These findings indicate that fCrd is inactive as isolated and requires preincubation with NADH for activation. That the length of the time lag of anaerobic cinnamate reduction started by fCrd did not depend on fCrd concentration (data not shown) indicated that the activation is not associated with changes in the oligomeric structure. Furthermore, the red trace in Fig. 6A demonstrates that the activation is much faster in the absence of cinnamate than in its presence (blue curve). The duration of Crd preincubation with NADH for the red curve in Fig. 6A was approximately 3 min and this time was thus sufficient for nearly complete activation in the absence of the electron acceptor. Preincubation with 5 mM dithionite for 3 min before starting the reaction by NADH activated fCrd to the same level (data not shown). These findings suggested that the activation is likely caused by the reduction of a prosthetic group in fCrd rather than by mere NADH binding to it.

For further analysis, the product formation curve for the reaction started by Crd in Fig. 6A (blue curve) was converted into the first derivative (i.e., specific activity) plot (Fig. 6B), and its sigmoidal shape clearly indicated that the activation process is not a one-step reaction. Fitting experiments demonstrated a poor fit of the kinetic scheme involving two consecutive steps (red line) but a reasonably good fit for a scheme involving three such steps (green line). In these fittings, we assumed that the rate constants for each step are identical because this relationship corresponds to the largest possible time lag of the final product formation. The fitted value of the rate constant was 0.060 ± 0.003 s^-1^ for the three-step reaction. Three is thus the minimal number of the steps involved in slow Crd activation by NADH.

A similar slow activation of Crd was observed in the acrylate reduction started by NADH (Fig. 6C). The maximal activity measured with this substrate (3.5 s^-1^) was 23% of that with cinnamate. For comparison, this activity was only 1.3 s^-1^ with MV as the electron donor (6% of the activity with cinnamate). A similar kinetic analysis indicated that Crd activation by NADH in the presence of acrylate involves at least two steps (Fig. S1), both with the rate constant of 0.27 ± 0.02 s^-1^. Using a low dead-time cuvette for kinetic measurements, we directly demonstrated that Crd activation by NADH in the absence of acrylate (Fig. S1, left panel, squares) was faster than in its presence (Fig. S1, left panel, circles). A similarly kinetic analysis have revealed that the activation also involved two consecutive steps with the average rate constant of 0.75 ± 0.15 s^-1^.

If MV was used as the electron donor with various substrates (Table 1), the progress curves were always linear irrespective of the order of fCrd and substrate additions. This result suggested that the activation/deactivation of fCrd is somehow associated with its NADH-binding *NADH:flavin* domain. This hypothesis was supported by the observation that the NADH:FMN reduction by fCrd also demonstrated slow activation if the reaction was started by NADH (Fig. 6D). The activation kinetics was more complex in this case and demonstrated a poor fit of the consecutive three-step model (Fig. S1).

Crd activated by preincubation with NADH remained active until all NADH was consumed in the enzymatic reaction, but slowly returned to the inactive state afterwards. In the experiment illustrated in Fig. 7A, the reaction was started by adding acrylate to preactivated fCrd and proceeded linearly until NADH exhaustion. A new portion of NADH restored Crd activity to the level that depended on the time interval *t* before the second NADH addition (Fig. 7B). The time-courses became nonlinear at high *t*, indicating activation as in Fig. 6C (blue trace) to a final rate similar to those in Figs 6C (red trace) and 7A (left trace). These findings indicate that active fCrd is gradually deactivated when depleted of NADH but time-dependently restores full activity after NADH addition. fCrd deactivation was a first-order reaction with a half-time of 4.7 min (Fig. 7B).

**Figure 7.**
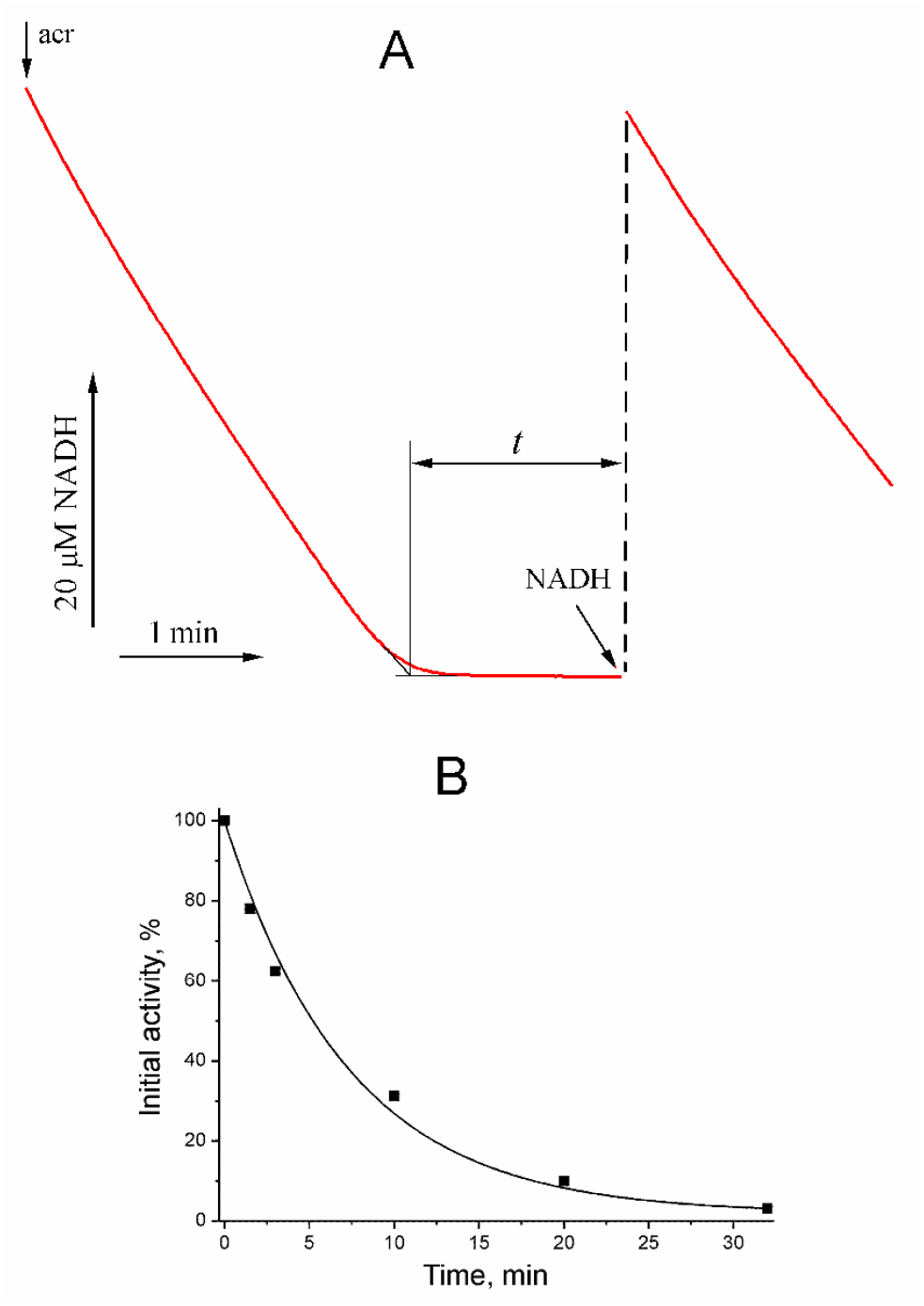
(A) NADH oxidation by fCrd in the presence of 1 mM acrylate (acr). When the reaction ended because of NADH consumption, its concentration was restored to the initial level after a 1.5-min pause. Conditions were as for Fig. 6C. (B) The initial activity of NADH conversion measured for the second portion of NADH as a function of the time interval *t* before its re-addition. 100% refers to the initial activity estimated from the left curve in panel A. The theoretical curve is for the first-order reaction with a half-time of 4.7 min.

Under aerobic conditions, the NADH-oxidizing activity of Crd was extremely low and did not increase with time (Fig. 6A, black trace). This observation contrasts the data reported for other NADH:α,β-enoate and NADH:flavin reductases, which readily oxidize NADH with O_2_ as the main electron acceptor (Bertsova et al., 2020; 2021; Hertzberger et al., 2014).

Interestingly, NADPH demonstrated only low efficiency as an electron donor compared with NADH. In the corresponding experiment, fCrd preactivated by incubation with 30 μM NADH was allowed to convert acrylate until all NADH was consumed and immediately supplemented with 120 μM NADPH. The measured rate of NADPH oxidation was 35 times lower than the rate of NADH oxidation. This makes NADH the most likely physiological donor of the reducing equivalents for Crd.

### 2.4 The roles of different domains in the catalytic activities of Crd

Crd is thus capable of catalyzing reactions between different natural and artificial donors and acceptors of electrons. To determine which Crd domains are responsible for different reactions, we measured the activities of the apo- and flavinylated forms of Crd and its deletion variants. The MV:cinnamate reductase activity was high with fCrd and much lower with aCrd (no covalently bound FMN) and d3 (only *FAD binding 2* domain) (Table 2). These data indicated that, like in *Klebsiella pneumoniae* NADH:fumarate reductase (Bertsova et al., 2020), the *FAD binding 2* domain is required for cinnamate reduction and the electron pathway from MV to this domain involves the covalently bound FMN of the *FMN bind* domain.

**Table 2.**
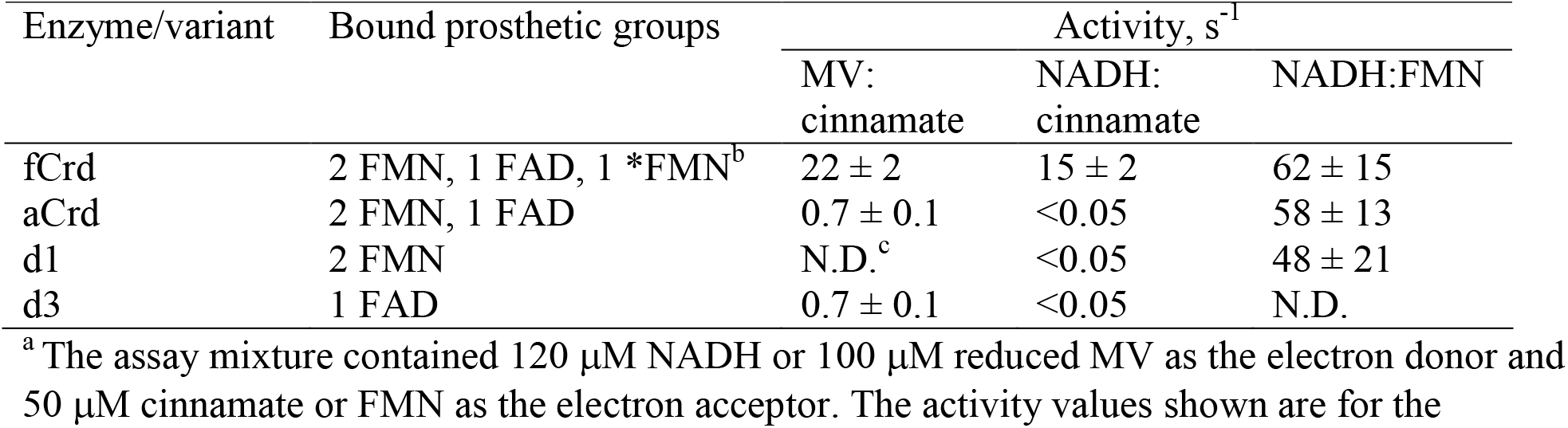
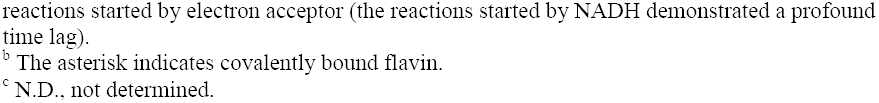
The catalytic activities of Crd and its deletion variants^a^

| Enzyme/variant | Bound prosthetic groups | Activity, s<sup>-1</sup> |  |  |
| --- | --- | --- | --- | --- |
|  |  | MV:<br>cinnamate | NADH:<br>cinnamate | NADH:FMN |
| fCrd | 2 FMN, 1 FAD, 1 *FMN<sup>b</sup> | 22 ± 2 | 15 ± 2 | 62 ± 15 |
| aCrd | 2 FMN, 1 FAD | 0.7 ± 0.1 | <0.05 | 58 ± 13 |
| d1 | 2 FMN | N.D.<sup>c</sup> | <0.05 | 48 ± 21 |
| d3 | 1 FAD | 0.7 ± 0.1 | <0.05 | N.D. |

<sup>a</sup> The assay mixture contained 120 μM NADH or 100 μM reduced MV as the electron donor and 50 μM cinnamate or FMN as the electron acceptor. The activity values shown are for the
reactions started by electron acceptor (the reactions started by NADH demonstrated a profound time lag).
<sup>b</sup> The asterisk indicates covalently bound flavin.
<sup>c</sup> N.D., not determined.

In contrast, high NADH:FMN reductase activity was demonstrated not only by the full-size flavinylated enzyme (fCrd) but also by its apo-form and the d1 variant (only *NADH:flavin* domain). This activity is thus clearly catalyzed by the *NADH:flavin* domain. Noteworthy, all the NADH:FMN reductase activities developed slowly in time when the reaction was started by NADH but instantly when FMN was the last added component, as in Fig. 6D.

Full NADH:cinnamate activity was manifested by fCrd but not aCrd that lacks covalently bound FMN. These findings emphasized the role of the covalently bound FMN in electron transfer between the NADH dehydrogenase part (d1) and the cinnamate reductase part (d3) in Crd.

The separate *FAD binding 2* domain (d3 variant) demonstrated measurable activity (0.60 ± 0.05 s^-1^) in the FMN:cinnamate reduction, raising the possibility of catalyzing complete NADH:cinnamate reductase reaction by a 1:1 mixture of d1 and d3 in the presence of medium FMN. Such activity was indeed detected, and, although its value, 3.2 ± 0.4 s^-1^, was less than for fCrd, it significantly exceeded the FMN:cinnamate reductase activity of d3. This seemingly unexpected result could be, however, explained by different conditions used for the two reactions. The assay medium for the FMN:cinnamate reductase activity contained, beside reduced FMN, considerable amounts of its oxidized form, which could inhibit the reaction by decreasing the redox potential of the FMNH_2_^free^/FMN^free^ pair. In contrast, virtually all FMN was in the reduced form under steady-state conditions in the NADH:cinnamate reductase reaction catalyzed by d1+d3. By demonstrating full NADH:cinnamate reductase activity in the presence of medium FMN, the d1 and d3 mixture resembles yeast soluble fumarate reductase (Kim et al., 2018). In the latter enzyme, the roles of d1 and d3 are played by separate enzymes, with free cytoplasmic flavins acting as electron carriers between them.

### 2.5 The three-dimensional structure of Crd

The 3D structure of heterodimeric Crd was predicted using AlphaFold2 (Jumper et al., 2021). The Ramachandran plot indicated that 96.7% residues have favorable dihedral angles in this structure. It can be divided into two parts connected by a flexible linker (Fig. 8). One of them, corresponding to d1 fragment, is formed by CrdA and the homologous N-terminal *NADH:flavin* domain of CrdB and resembles homodimeric NADH:flavin reductases (Agarwal et al., 2006; Koike et al., 1998). The other part contains two other CrdB domains—*FAD binding 2* and *FMN bind*. The predicted structure of the CrdB subunit is similar to but not identical with that earlier predicted using the previous software version (AlphaFold) and found in UniProt. The major differences are seen in the *FAD binding 2* domain; furthermore, the latter structure does not contain CrdA.

**Figure 8.**
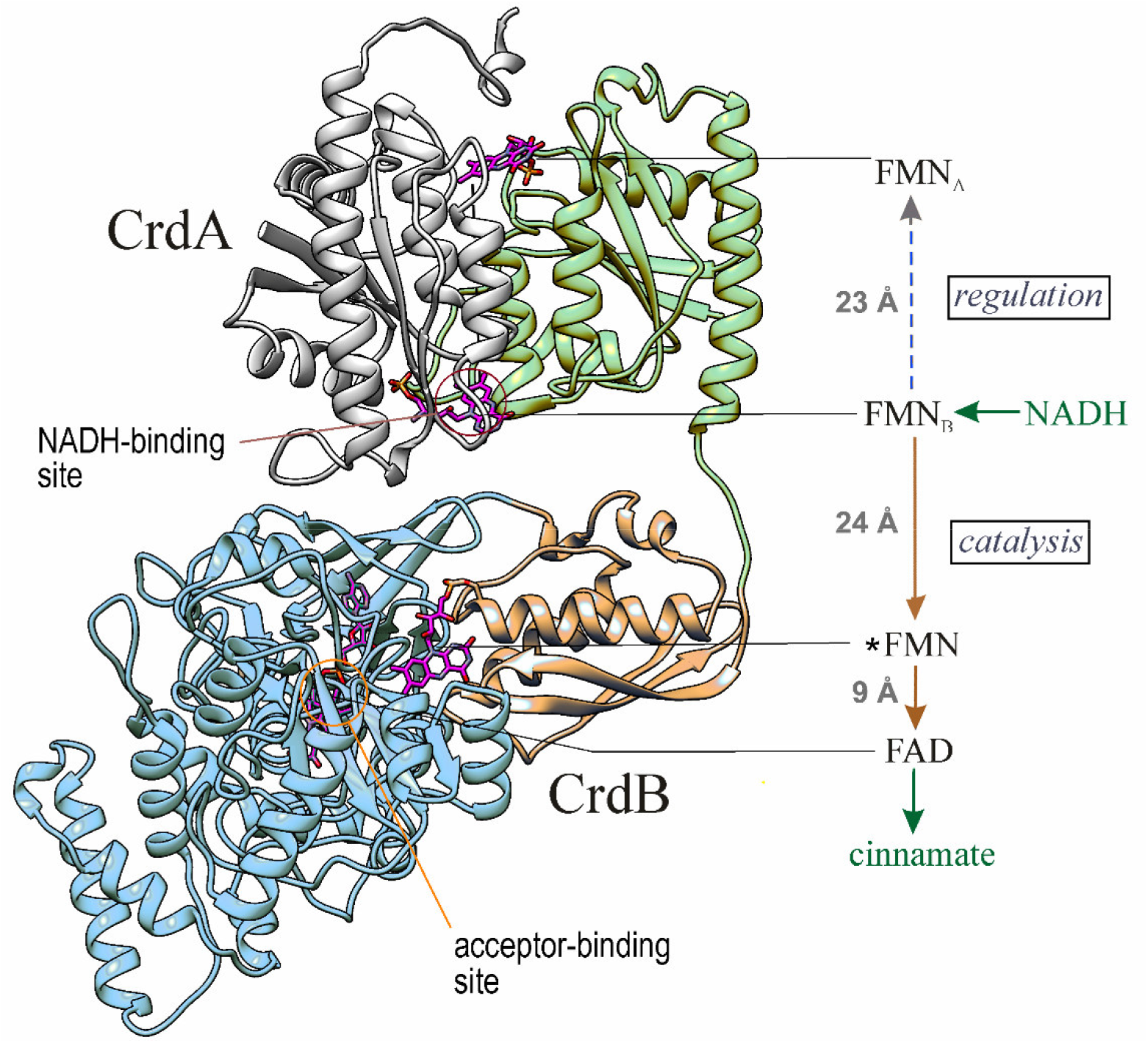
The predicted three-dimensional structure of Crd with bound FAD and three FMNs in a cartoon representation. The domains are colored like in Fig. 1. The right part shows schematically the edge-to-edge distances between the prosthetic groups and the proposed pathways for electron and hydride ion transfer (indicated by red and green arrows, respectively).

Four flavin groups (three FMNs and one FAD in their oxidized states) and one each of NADH and cinnamate molecules were docked into the modelled structure using AutoDock Vina (Trott et al., 2010) in the following sequence: *FMN, FAD, FMN_B_, FMN_A_, NADH, cinnamate. The locations of the docked flavin groups (Fig. 8) were similar to those in the experimentally determined structures of NAD(P)H:FMN reductase (PDB ID: 1×77; (Agarwal et al., 2006)), the NqrC subunit of Na^+^-translocating NADH:quinone oxidoreductase (PDB ID: 4XA7; (Borshchevskiy et al., 2015)), and urocanate reductase (PDB ID: 6T87; (Venskutonytė et al., 2021)) (Fig. 8). These three proteins are homologous to the *NADH:flavin, FMN bind*, and *FAD binding 2* domains of Crd, respectively.

The d1 part of Crd contains two potential pseudo-symmetrical NADH-binding sites near FMN_A_ and FMN_B_, similar to those found in the homodimeric NADH:flavin reductase (Koike et al., 1998). However, only the FMN_B_-adjacent site could be occupied by NADH because the entrance to the FMN_A_-adjacent site is blocked by the C-terminal segment of CrdB (Fig. S2). Instead, the positions of the docked NADH molecule concentrated in two areas of the protein near FMN_A_. The shortest distance between NADH and FMN_A_ was 10 Å, which is too large for the hydride ion transfer between them (for comparison, the distance between NADH and FMN_B_ was only 4 Å). Similar docking experiments provided no evidence for a competent NADH-binding site in the middle, *FMN bind* domain of CrdB. The location of docked cinnamate in the *FAD binding 2* domain of CrdB was similar to that of urocanate in urocanate reductase (Venskutonytė et al., 2021).

The edge-to-edge distance between FAD and *FMN in the resulting model is approximately 9 Å (Fig. 8), allowing rapid electron transfer between these flavin groups, consistent with the finding that this step is not rate-limiting in NADH:α,β-enoate reductases (Bertsova et al., 2020). In contrast, the FMN_B_ and *FMN groups are separated by 24 Å in the model (Fig. 8), a distance at which rapid electron transfer is not feasible. Clearly, Crd should adopt a different conformation to make the FMN_B_ and *FMN groups closer to each other and allow electron transfer between them in the catalytic reaction. Interestingly, the same step is rate-limiting in the electron transfer in canonical NADH:α,β-enoate reductases (Bertsova et al., 2020), indicating that the above distance is similarly non-optimal in these enzymes.

The validity of the model predicted by AlphaFold2 with the docked flavins was confirmed by MD simulations (Fig. S3). Specifically, none of the four flavin groups was lost during the simulations. Furthermore, the simulations did not change significantly the structure predicted by AlphaFold2, except for a rigid-body movement of the d1 part, connected by a flexible linker, by approximately 11 Å relative the rest of the molecule (Fig. S3A,C). This result showed that Crd structure is indeed flexible. However, the flavin-to-flavin distances were not appreciably different in the simulated structure (Fig. S3B) and remained virtually unchanged in the analogous structure with the flavins in their reduced forms (data not shown).

### 2.6 Induction of Crd synthesis in *V. ruber* cells by unsaturated carbonic acids

The effects of cinnamate and other unsaturated carbonic acids added to *V. ruber* growth medium on the MV:cinnamate reductase activity in cleared cell lysates were measured. All cultivations were performed in duplicate both in the presence and in the absence of O_2_.

As Table 3 makes clear, cinnamate added at its maximal concentration (10 mM) that did not yet suppress cell growth caused appreciable induction of Crd synthesis. The effect was observed under both aerobic and anaerobic conditions but was maximal in the former case.

**Table 3.** Correlation between the MV:cinnamate reductase activity and *crd*B transcription level in the *V. ruber* cells cultivated in different conditions.

| Growth conditions | Crd activity (nmol·min<sup>-1</sup>·mg<sup>-1</sup>) | *crdB* mRNA/rRNA × 10<sup>-6</sup> |
| --- | --- | --- |
| +O<sub>2</sub>, no cinnamate | 0.5 ± 0.2 | 0.8 ± 0.1 |
| +O<sub>2</sub>, 10 mM cinnamate | 19 ± 3 | 25 ± 2 |
| -O<sub>2</sub>, no cinnamate | 1.2 ± 0.4 | 2.1 ± 0.2 |
| -O<sub>2</sub>, 10 mM cinnamate | 86 ± 11 | 31 ± 4 |

Parallel measurements indicated a more than ten-fold increase in the gene *crd*B transcription by cinnamate, providing further support for assigning the MV:cinnamate reductase activity to Crd in *V. ruber* cells.

The other α,β-unsaturated carbonic acids acting as Crd substrates, except for acrylate, could induce Crd synthesis under anaerobic conditions (Fig. 9). These compounds were added at 2 mM concentrations because the cells did not grow anaerobically at higher concentrations of acrylate and caffeate. Noteworthy, Crd induction by cinnamate was not maximal at this concentration, based on the activity values in Table 3 and Fig. 9.

**Figure 9.**
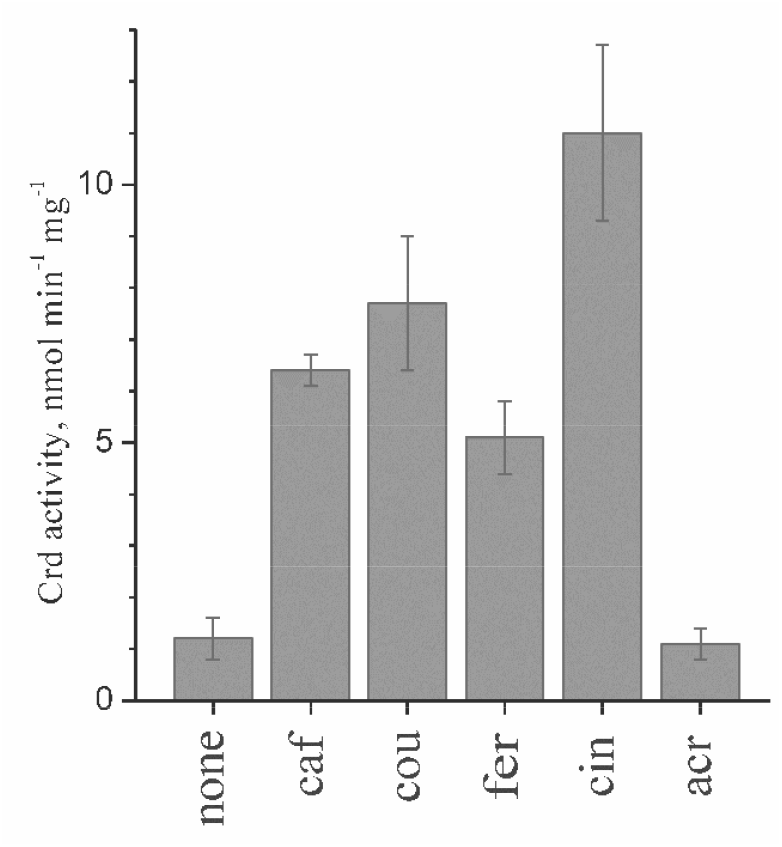
MV:cinnamate reductase activity of the *V. ruber* cells grown anaerobically in the presence of 2 mM acrylate (acr), cinnamate (cin), *p*-coumarate (cou), caffeate (caf), or ferulate (fer). Error bars represent the standard deviations for two cultivations.

## 3 DUSCUSSION

While sharing overall structural similarity with other microbial three-domain NADH:2-enoate reductases, such as the fumarate reductase of *K. pneumoniae* and Kinetoplastida and acrylate reductase of *V. harveyi*, the *V. ruber* cinnamate reductase Crd displays significant differences. In all these enzymes, α,β-enoate is reduced in *FAD binding 2* domain by the electrons transmitted via a covalently bound FMN residue of *FMN bind* domain (Bertsova et al., 2020). However, the NADH-oxidizing activity belongs to *NADH:flavin* domain in Crd but to either *OYE*-*like* domain (in *K. pneumoniae* NADH:fumarate and *V. harveyi* NADH:acrylate reductases) or *FAD binding 6* domain (in Kinetoplastida NADH:fumarate reductases).

Furthermore, only Crd contains an additional NADH-reducing subunit (CrdA) formed by a single *NADH:flavin* domain, which is homologous to the analogous domain of CrdB (51% identity, 65 % similarity). Its gene (*crd*A) flanks the *crd*B gene in one chromosome (Fig. 1C), which, together with the finding that the tag-less CrdA co-purifies with 6×His-tagged CrdB on a Ni-column, provides strong evidence that Crd functions as a tight CrdA/CrdB complex *in vivo*. The observed noncovalently bound FMN:FAD stoichiometry of 2:1 is fully consistent with the heterodimeric structure of Crd. Noteworthy, prokaryotic NADH:flavin reductases are generally encoded by single genes but function as homodimers containing two bound FMN molecules at the dimer interface (Agarwal et al., 2006; Koike et al., 1998). The homodimeric structure apparently prevents direct fusion of the NADH:flavin reductase domain with other functional domains to form an extended multidomain polypeptide.

The presence of the *NADH:flavin* domain in Crd instead of the *OYE*-*like* or *FAD binding 6* domain appears to provide several advantages. The alternative three-domain NADH:2-enoate reductases, containing the latter domains, additionally catalyze O_2_ reduction under aerobic conditions, forming reactive oxygen species (Bertsova et al., 2020; Wargnies et al., 2021). This additional activity may be harmless for strictly anaerobic bacteria but may cause cell damage in facultatively anaerobic microorganisms, like *V. ruber*, when they meet O_2_. Remarkably, reactive oxygen species formation in *V. ruber* by Crd is prevented by a unique regulation mechanism which reversibly switches off Crd activity at high O_2_ concentrations (Fig. 6A). The transition between the active and inactive states appears to be controlled in Crd by the redox state of one or both FMN molecules in the *NADH:flavin* domains. This conclusion is supported by the observations that Crd could be activated by both NADH and dithionite and that the d1 but not d3 fragment demonstrated similar time-dependent activation of its partial enzymatic activity.

Crd regulation appears to be associated with the heterodimeric structure of the NADH dehydrogenase part, formed by CrdA and the homologous N-termninal domain of CrdB (Fig. 8). This part of Crd resembles NADH:flavin reductases (Agarwal et al., 2006; Koike et al., 1998) and similarly contains two noncovalently bound FMN groups and two potential sites of NADH oxidation. However, the competence of the putative NADH-binding site near FMN_A_ seems questionable because of the unfavorable steric effect of the C-terminal loop of CrdA. While electron transfer between FMN_B_ and FMN* is perceivable assuming FMN* and/or *NADH:flavin* dimer flexibility, the high distance between FMN_B_ and FMN_A_ (Fig. 8) can hardly decrease similarly. Hence, the electron transfer between these flavin groups should be always much slower than enzyme turnover (Page et al., 1999). These structural considerations suggest that FMN_B_ and FMN_A_ are good candidates for the catalytic and regulating groups, respectively. That these flavins have distinct roles is consistent with significant differences in the amino acid sequences of the two *NADH:flavin* domains of Crd, contrasting the case of homodimeric and, hence, symmetrical NADH:flavin reductases in which two FMN sites are identical (Koike et al., 1998). One can further speculate that Crd activation involves the stepwise reduction of FMN_B_ and FMN_A_ causing a conformational change that forms a functional electron transfer chain linking the donor and acceptor.

That the electron acceptors slow down Crd activation by NADH raises the possibility that the bound acceptor competes with the regulating prosthetic group, likely FMN_A_, for the electrons coming from NADH. In this case, the activation rate should inversely correlate with the rate of acceptor reduction. Such a correlation was indeed observed—cinnamate was reduced faster (Table 1) and decelerated Crd activation more compared with acrylate (Fig. 6). In this context, the lack of Crd activation under aerobic conditions may mean that O_2_ can oxidize the regulating prosthetic group directly, without using the long electron transfer chain involved in the case of cinnamate and acrylate. An alternative explanation, suggested by the dependence of activation kinetics on cinnamate concentration, is that the bound acceptor sterically prevents Crd interaction with NADH.

Despite the similarity of the acceptor-binding amino acid residues in the *FAD binding 2* domains of acrylate reductase and Crd (Fig. 1B), the latter enzyme exhibits a 1,000–6,000-fold lower catalytic efficiency against acrylate than against cinnamate and its derivatives (Table 1). Furthermore, (hydroxy)cinnamic acids, but not acrylate, induced Crd synthesis in *V. ruber* cells. Clearly, the positions of amino acid residues interacting with the electron-acceptor carboxylates in fumarate reductases are not the only determinants of the substrate specificity in other NADH:2-enoate reductases. The reduction of cinnamate and its derivatives, toxic to cells, is thus the most likely physiological function of Crd, and this inference is supported by their induction of Crd synthesis in both anaerobic and aerobic conditions (Table 3). Additionally, cinnamate and *p*-coumarate, being the products of phenylalanine and tyrosine deamination, may support anaerobic respiration of *V. ruber* during Stickland metabolism (Liu et al., 2022). Two other “good” substrates of Crd, caffeate and ferulate, are the precursors of lignin and various phenolic secondary metabolites in plants. That is why these compounds are abundant in terrestrial niches, soil in particular, but are relatively rare in the marine habitats, where they are only synthesized in some algae (Fernando et al., 2016; Martone et al., 2009).

Crd-like reductases are relatively rare among marine microorganisms, seemingly, because (hydroxyl)cinnamic acids are underrepresented in the habitat. BLAST search has revealed Crd-like proteins only in the following marine bacteria: *Vibrio gazogenes, Vibrio salinus, Vibrio spartinae*, and *Vibrio tritonius*. However, such reductases are abundant among anaerobic terrestrial bacteria (the *Clostridium, Klebsiella, Citrobacter, Aeromonas, Paenibacillus, Streptococcus*, and many other genera), including those forming human intestinal microbiome. The presence of substantial amounts of phenylpropionate and *p*-hydroxyphenylpropionate in the mammalian intestine has long been known (Smith et al., 1996) but their production was attributed to a different enzyme, bacterial two-domain 2-enoate reductase (Tischer et al., 1979). The wide distribution of Crd-like enzymes among intestinal bacteria suggests that they are also involved in phenylpropionate and *p*-hydroxyphenylpropionate formation in the mammalian intestine.

Two-domain 2-enoate reductases are industrial biocatalysts increasingly used to produce chiral intermediates, pharmaceuticals and agrochemicals (Knaus et al., 2016; Mordaka et al., 2018). A disadvantage of these enzymes is that they are readily and irreversibly inactivated by oxygen (Mordaka et al., 2018; Tischer et al., 1979). The lower sensitivity of Crd to oxygen makes it a promising alternative for industrial application.

## 4 EXPERIMENTAL PROCEDURES

### 4.1 Bacterial strains and growth conditions

*V. ruber* DSM 16370 was obtained from the Leibniz Institute collection of microorganisms and cell cultures (DSMZ). The cells were grown aerobically or anaerobically at 28 °C in a mineral medium containing 20 g/L NaCl, 0.75 g/L KCl, 1.2 g/L MgSO_4_×7(H_2_O), 0.5 g/L NH_4_Cl, 0.5 mM Na_2_HPO_4_, 2 g/L sucrose, 0.5 g/L yeast extract, and 50 mM Tris-HCl (pH 8.0). When required, the medium was supplemented with 2–10 mM acrylate, cinnamate, *p*-coumarate, caffeate, or ferulate. Cleared cell lysate was prepared as described elsewhere (Bertsova et al., 2019) and assayed for MV:cinnamate reductase activity. *E. coli* cells were grown at 37 °C in LB. Where indicated, the LB medium was supplemented with 20 μg/mL chloramphenicol and (or) 100 μg/mL ampicillin.

### 4.2 Construction of expression vectors

The expression vector for the CrdAB protein with C-terminal 6×His-tagged CrdB was constructed by amplifying the *crd*-operon from the genomic *V. ruber* DNA. A high-fidelity Tersus polymerase (Evrogen) and the primers 5’-GGGACTCTGCTGCCTTGCT / 5’-CACTGCTTGGTCCATTTCACT were used. The resulting 3276-bp fragment was cloned into the pBAD-TOPO vector (Invitrogen), yielding the pTB_CRD4 plasmid. The plasmid was transformed into *E. coli* / pΔhis3 (Bertsova et al., 2013) or BL21 strains.

The expression vector for the d1 fragment (full-size CrdA and residues 1–184 of CrdB) was constructed by amplifying the corresponding DNA fragment of the *crd*-operon by PCR with a Tersus PCR kit and the primers 5’-GGGACTCTGCTGCCTTGCT / 5’-CTCTACATTCAGTTGGTTTGAGAT using *V. ruber* genomic DNA as the template. The resulting 1401-bp fragment was cloned into the pBAD-TOPO vector, yielding the pTB_d1CRD17 plasmid. To construct an expression vector for the d3 fragment (residues 281– 806) of CrdB, the 1688-bp fragment was amplified by PCR with the 5’-<u>CATATG</u>CTGAAGAAACGGCCCAAAC / 5’-<u>CTCGAG</u>ACTCAATGGTGATGGTG primer pair (restriction sites for *Nde*I and *Xho*I are underlined) and the pTB_CRD4 plasmid as the template. The amplified fragment was cloned into the pSCodon vector (Delphi Genetics) using the *Nde*I/*Xho*I sites, resulting in the plasmid pSC_d3CRD2. The pTB_d1CRD17 and pSC_d3CRD2 plasmids were transformed into *E. coli* BL21 strain.

### 4.3 Crd production

The 6×His-tagged Crd proteins with or without covalently bound FMN (fCrd and aCrd, respectively) and the d1 and d3 fragments of Crd were produced in *E. coli* BL21 or /pΔhis3 cells and purified using metal chelate chromatography as described previously (Bertsova et al., 2014). The extinction coefficients ε_450nm_ for fCrd, aCrd, d1 fragment, and d3 fragment determined after SDS treatment (Bertsova et al., 2022) were 42, 31, 22, and 11 mM^-1^ cm^-1^, respectively.

### 4.4 Analysis of flavins

Non-covalently bound flavins were extracted from Crd preparations by trifluoroacetic acid (TFA) and separated by HPLC (Bertsova et al., 2014). SDS-PAGE was performed using 12.5% (w/v) polyacrylamide gels (Laemmli, 1970). The gels were stained for protein with PageBlue™ solution (Fermentas). Covalently bound flavins were detected by scanning unstained gels with a Typhoon™ FLA 9500 laser scanner (GE Healthcare) with excitation at 473 nm and the detection of emission using the SYBR Green II protocol according to the manufacturer’s recommendations.

For mass-spectral measurements, gel pieces of approximately 2 mm^3^ were excised from the protein bands of the Coomassie-stained gel and destained by incubating in two 0.1-mL volumes of 40% (by volume) acetonitrile solution containing 20 mM NH_4_HCO_3_, pH 7.5, dehydrated with 0.2 mL of 100% acetonitrile and rehydrated with 5 μL of the digestion solution containing 15 μg/mL sequencing-grade trypsin (Promega) in 20 mM aqueous solution of NH_4_HCO_3_, pH 7.5. Digestion was carried out at 37°C for 1 h. The resulting peptides were extracted with 5 μL 30% acetonitrile solution containing 0.5% TFA. A 1-μL aliquot of the in-gel tryptic digest extract was mixed with 0.5 μL of 2,5-dihydroxybenzoic acid solution (40 mg/mL) in 30% acetonitrile containing 0.5% TFA and left to dry on a stainless-steel target plate. MALDI-TOF MS analysis was performed on an UltrafleX?treme MALDI-TOF-TOF mass spectrometer (Bruker Daltonik, Germany). The MH^+^ molecular ions were measured in a reflector mode; the accuracy of monoisotopic mass peak measurement was within 30 ppm. Spectra of fragmentation were obtained in a LIFT mode, the accuracy of daughter ion measurement was within 1 Da. Mass-spectra were processed with the FlexAnalysis 3.2 software (Bruker Daltonik, Germany). Protein identification was carried out by MS+MS/MS ion search, using Mascot software version 2.3.02 (Matrix Science), through the Home NCBI Protein Database. One each of missed cleavage, Met oxidation, Cys-propionamide, and Ser(Thr)-FMN were permitted. Protein scores greater than 94 were considered significant (p < 0.05).

### 4.5 Enzymatic activities

NADH- and methyl viologen (MV)-supported activities of Crd preparations were determined at 25 °C by following NADH or MV oxidation spectrophotometrically at 340 or 606 nm, respectively (Bertsova et al., 2020). The assay medium contained 1 mM MV, 1 mM electron acceptor, and 100 mM Tris-HCl (pH 8.0) or 150 μM NADH, 0.05–1 mM electron acceptor, 10 mM glucose, 5 U/mL glucose oxidase, 5 U/mL catalase, and 100 mM Mes-KOH (pH 6.5) in a completely filled and sealed 3.2-mL cuvette. MV was pre-reduced with sodium dithionite until the absorbance at 606 nm of approximately 1.5 was obtained, which corresponds to the formation of ∼100 μM reduced MV. Unless otherwise noted, Crd or its variant was preincubated with NADH or MV for 2–3 min before the reaction was started by adding an electron acceptor. The NADH-supported reaction was also measured under aerobic conditions without glucose oxidase and catalase. Crd activities were calculated using the experimentally verified NADH:acceptor and MV:acceptor stoichiometries of 1:1 and 2:1, respectively.

The assay medium for measuring FMNH_2_:cinnamate reductase activity contained 100 μM FMN, 1 mM cinnamate, and 100 mM Tris-HCl (pH 8.0). FMN was pre-reduced with sodium dithionite until the absorbance at 450 nm of approximately 0.3 was obtained, which corresponds to formation of ∼70 μM reduced FMN. FMNH_2_ oxidation was followed spectrophotometrically at 450 nm.

Phenylpropionate dehydrogenase activity was assayed by measuring the reduction of 2,6-dichlorophenolindophenol (DCPIP, ε_600_ = 22 mM^-1^ cm^-1^) at 600 nm (Ells, 1959). The assay mixture contained 2 mM Tris-phenylpropionate, 2 mM phenazine methosulphate, 25 μM DCPIP, and 100 mM Mes-KOH (pH 6.5).

The Michaelis-Menten parameters of MV-supported acrylate reduction by Crd were estimated from duplicate measurements of the initial rates of MV reduction using 0.03–10 mM acrylate and 40 nM Crd concentrations. For other electron acceptors, rates were obtained from the integral kinetics of their reduction until its completion, as monitored by absorbance at 606 nm. Rates were estimated at 100 time points along the progress curve as the slopes of the tangents (-*d*[MV]/*d*t) using MATLAB (The MathWorks, Inc.). The residual electron acceptor concentration was calculated at each point from *A*_606_ using the MV:acceptor stoichiometry of 2:1 and assuming that the limiting value of *A*_606_ corresponds to 100% conversion of the electron acceptor. The Michaelis-Menten equation was fitted to the rate data using non-linear regression analysis.

### 4.6 Identification of the product of cinnamate reduction

Cinnamate reduction was performed in the medium containing 100 μM cinnamate, 10 mM sodium dithionite, 50 μM MV, and 100 mM Tris-HCl (pH 8.0). The reaction was initiated by adding 0.5 μM Crd and terminated after 30 min by adding 5% (vol./vol.) HClO_4_. In the control experiment, HClO_4_ was added before Crd. Precipitated protein was removed by centrifugation, and the substrate and products were extracted from the samples with diethyl ether (2 × 1.5 mL). The ether was evaporated under a stream of air, and the residue was dissolved in 1 mL of 25 mM potassium phosphate (pH 6.5) (medium A). The samples were separated by HPLC on a ProntoSil-120-5-C18 AQ column using a Milichrom A-02 chromatograph (both from Econova, Novosibirsk, Russia). The column was pre-equilibrated with medium A and eluted with a linear gradient from 0% to 10% methanol in medium A at a flow rate of 0.2 mL/min with UV detection at 210 and 258 nm.

### 4.7 Quantitative reverse transcription polymerase chain reaction (RT-qPCR)

RNA extraction from *V. ruber* cells and cDNA synthesis were performed as described previously (Bertsova et al., 2019). RT-qPCR assays were performed with qPCRmix-HS SYBR kit (Evrogen), using the cDNA preparations as templates and 5’-GCAAGCGCATTTCAAGGAC / 5’-CCAGCGGGTGAAGTTCGTAG as primer pair for *crd*B. 16S rRNA was used for data normalization (the primer pair 5’-CAGCCACACTGGAACTGAGA / 5’-GTTAGCCGGTGCTTCTTC). Serial dilutions of *V. ruber* genomic DNA, which contains the genes for CrdB and 16S rRNA in a 1:8 ratio, were used for calibration.

### 4.8 Bioinformatics

The three-dimensional structure of flavin-deficient Crd was predicted from the amino acid sequences of CrdA and CrdB using AlphaFold2 (version 2.2.0) (Jumper et al., 2021) through the ColabFold advanced notebook (Mirdita et al., 2022). Multimer model prediction was performed using default parameter settings.

Ligands (flavins, NADH, cinnamate) were docked in sequence into the obtained Crd structure using AutoDock Vina (Trott et al., 2010). Space cells ranging from 12×12×12 to 30×30×30Å^3^ were selected for docking in the appropriate regions of the protein based on the known structures of homologous proteins. Each docking experiment was performed in triplicate. The derived Crd structure containing one FAD and three FMN groups was finally equilibrated by molecular dynamics (MD) simulations for 150 ns with AMBER 22 package (Case et al., 2022) (http://ambermd.org/). Details of the simulation are found in Supplement (Fig. S3 legend).

## AUTHOR CONTRIBUTIONS

AVB – the conception of the study; YVB, MVS, VAA, AAB, and AVB – the acquisition and analysis of the data; AAB and AVB – writing of the manuscript.

## ACKNOWLEDGEMENTS

This work was supported by the Russian Science Foundation (project # 22-24-00133). MALDI MS and laser scanner facilities became available to us in the framework of the Moscow State University Development Program PNG 5.13.

## ETHICS STATEMENT

All strains, plasmids and primers will be made available upon reasonable request.

## REFERENCES

Agarwal, R., Bonanno, J.B., Burley, S.K., & Swaminathan, S. (2006). Structure determination of an FMN reductase from Pseudomonas aeruginosa PA01 using sulfur anomalous signal. Acta Crystallographica Section D, 62, 383–391. doi: 10.1107/S0907444906001600

Bertsova, Y.V., Fadeeva, M.S., Kostyrko, V.A., Serebryakova, M.V., Baykov, A.A., & Bogachev, A.V. (2013). Alternative pyrimidine biosynthesis protein ApbE is a flavin transferase catalyzing covalent attachment of FMN to a threonine residue in bacterial flavoproteins. Journal of Biological Chemistry, 288, 14276–14286. doi: 10.1074/jbc.M113.455402

Bertsova, Y.V., Kostyrko, V.A., Baykov, A.A., & Bogachev, A.V. (2014). Localization-controlled specificity of FAD:threonine flavin transferases in Klebsiella pneumoniae and its implications for the mechanism of Na+-translocating NADH:quinone oxidoreductase. Biochimica et Biophysica Acta, 1837, 1122–1129. doi: 10.1016/j.bbabio.2013.12.006

Bertsova, Y.V., Kulik, L.V., Mamedov, M.D., Baykov, A.A., & Bogachev, A.V. (2019). Flavodoxin with an air-stable flavin semiquinone in a green sulfur bacterium. Photosynthesis Research, 142, 127–136. doi: 10.1007/s11120-019-00658-1

Bertsova, Y.V., Oleynikov, I.P., & Bogachev, A.V. (2020) A new water-soluble bacterial NADH: fumarate oxidoreductase. FEMS Microbiology Letters, 367, fnaa175. doi: 10.1093/femsle/fnaa175

Bertsova, Y.V., Serebryakova, M.V., Baykov, A.A., & Bogachev, A.V. (2021). The flavin transferase ApbE flavinylates the ferredoxin:NAD+-oxidoreductase Rnf required for N2 fixation in Azotobacter vinelandii. FEMS Microbiology Letters, 368, fnab130. doi: 10.1093/femsle/fnab130

Bertsova, Y.V., Serebryakova, M.V., Baykov, A.A., & Bogachev, A.V. (2022). A novel, NADH-dependent acrylate reductase in Vibrio harveyi. Applied and Environmental Microbiology, 88, e0051922. doi: 10.1128/aem.00519-22

Besteiro, S., Biran, M., Biteau, N., Coustou, V., Baltz, T., Canioni, P., & Bringaud, F. (2002). Succinate secreted by Trypanosoma brucei is produced by a novel and unique glycosomal enzyme, NADH-dependent fumarate reductase. Journal of Biological Chemistry, 277, 38001–38012. doi: 10.1074/jbc.M201759200

Bogachev, A.V., Bertsova, Y.V., Bloch, D.A., & Verkhovsky, M.I. (2012). Urocanate reductase: Identification of a novel anaerobic respiratory pathway in Shewanella oneidensis MR-1. Molecular Microbiology, 86, 1452–1463. doi: 10.1111/mmi.12067

Bogachev, A.V., Baykov, A.A., & Bertsova, Y.V. (2018). Flavin transferase: the maturation factor of flavin-containing oxidoreductases. Biochemical Society Transactions, 46, 1161–1169. doi: 10.1042/BST20180524

Bolanos-Garcia, V.M., & Davies, O.R. (2006). Structural analysis and classification of native proteins from E. coli commonly co-purified by immobilised metal affinity chromatography. Biochimica et Biophysica Acta, 1760, 1304–1313. doi: 10.1016/j.bbagen.2006.03.027

Borshchevskiy, V., Round E., Bertsova, Y., Polovinkin, V., Gushchin, I., Ishchenko A, … Gordeliy, V. (2015). Structural and functional investigation of flavin binding center of the NqrC subunit of sodium-translocating NADH:quinone oxidoreductase from Vibrio harveyi. PLoS One, 10, e0118548. doi: 10.1371/journal.pone.0118548

Caldeira, J., Feicht, R., White, H., Teixeira, M., Moura, J.J., Simon, H., & Moura, I. (1996). EPR and Mössbauer spectroscopic studies on enoate reductase. Journal of Biological Chemistry, 271, 18743–18748. doi: 10.1074/jbc.271.31.18743

Case, D.A., Aktulga, H.M., Belfon, K., Ben-Shalom, I.Y., Berryman, J.T., Brozell, S.R., … Kollman, P.A. (2022). Amber 2022, University of California, San Francisco.

Ells, A.H. (1959). A colorimetric method for the assay of soluble succinic dehydogenase and pyridinenucleotide-linked dehydrogenases. Archives of Biochemistry and Biophysics, 85, 561–562. doi: 10.1016/0003-9861(59)90527-2

Fernando, I.P., Kim, M., Son, K.T., Jeong, Y., & Jeon, Y.J. (2016). Antioxidant activity of marine algal polyphenolic compounds: a mechanistic approach. Journal of Medicinal Food, 19, 615–628. doi: 10.1089/jmf.2016.3706

Giesel, H., & Simon, H. (1983). On the occurrence of enoate reductase and 2-oxo-carboxylate reductase in clostridia and some observations on the amino acid fermentation by Peptostreptococcus anaerobius. Archives of Microbiology, 135, 51–57. doi: 10.1007/BF00419482

Hertzberger, R., Arents, J., Dekker, H.L., Pridmore, R.D., Gysler, C., Kleerebezem, M., & de Mattos, M.J. (2014). H_2_O_2_ production in species of the Lactobacillus acidophilus group: a central role for a novel NADH-dependent flavin reductase. Applied and Environmental Microbiology, 80, 2229–2239. doi: 10.1128/AEM.04272213

Jumper, J., Evans, R., Pritzel, A., Green, T., Figurnov, M., Ronneberger, O., … Hassabis, D. (2021). Highly accurate protein structure prediction with AlphaFold. Nature, 596, 583–589. doi: 10.1038/s41586-021-03819-2

Kim, S., Kim, C.M., Son, Y.J., Choi, J.Y., Siegenthaler, R.K., Lee, Y., … Park, H.H. (2018). Molecular basis of maintaining an oxidizing environment under anaerobiosis by soluble fumarate reductase. Nature Communications, 9, 4867. doi: 10.1038/s41467-018-07285-9

Knaus, T., Toogood, H.S., & Scrutton, N.S. (2016). Ene-reductases and their applications. Green Biocatalysis. John Wiley & Sons, Inc. pp. 473–488.

Koike, H., Sasaki, H., Kobori, T., Zenno, S., Saigo, K., Murphy, M.E., … Tanokura, M. (1998). 1.8 Å crystal structure of the major NAD(P)H:FMN oxidoreductase of a bioluminescent bacterium, Vibrio fischeri: overall structure, cofactor and substrate-analog binding, and comparison with related flavoproteins. Journal of Molecular Biology, 280, 259–273. doi: 10.1006/jmbi.1998.1871

Kuno, S., Bacher, A., & Simon, H. (1985). Structure of enoate reductase from a Clostridium tyrobutyricum (C. spec. La1). Biological Chemistry Hoppe-Seyler, 366, 463–472. doi: 10.1515/bchm3.1985.366.1.463

Laemmli, U.K. (1970). Cleavage of structural proteins during the assembly of the head of bacteriophage T4. Nature, 227, 680–685. doi: 10.1038/227680a0

Light, S.H., Méheust, R., Ferrell, J.L., Cho, J., Deng, D., Agostoni, M., … Portnoy, D.A. (2019). Extracellular electron transfer powers flavinylated extracellular reductases in Gram-positive bacteria. Proceedings of the National Academy of Sciences of the United States of America, 116, 26892–26899. doi: 10.1073/pnas.1915678116

Liu, Y., Chen, H., Van Treuren, W., Hou, B.H., Higginbottom, S.K., & Dodd, D. (2022). Clostridium sporogenes uses reductive Stickland metabolism in the gut to generate ATP and produce circulating metabolites. Nature Microbiology, 7, 695–706. doi: 10.1038/s41564-022-01109-9

Martone, P.T., Estevez, J.M., Lu, F., Ruel, K., Denny, M.W., Somerville, C., & Ralph, J. (2009). Discovery of lignin in seaweed reveals convergent evolution of cell-wall architecture. Current Biology, 19, 169–175. doi: 10.1016/j.cub.2008.12.031

Mirdita, M., Schütze, K., Moriwaki, Y., Heo, L., Ovchinnikov, S., & Steinegger, M. (2022). ColabFold: making protein folding accessible to all. Nature Methods, 19, 679–682. doi: 10.1038/s41592-022-01488-1

Page, C.C., Moser, C.C., Chen, X., & Dutton, P.L. (1999). Natural engineering principles of electron tunnelling in biological oxidation-reduction. Nature, 402, 47–52. doi: 10.1038/46972

Serebryakova, M.V., Bertsova, Y.V., Sokolov, S.S., Kolesnikov, A.A., Baykov, A.A., & Bogachev, A.V. (2018). Catalytically important flavin linked through a phosphoester bond in a eukaryotic fumarate reductase. Biochimie, 149, 34–40. doi: 10.1016/j.biochi.2018.03.013

Shieh, W.Y., Chen, Y.W., Chaw, S.M., & Chiu, H.H. (2003). Vibrio ruber sp. nov., a red, facultatively anaerobic, marine bacterium isolated from sea water. International Journal of Systematic and Evolutionary Microbiology, 53, 479–484. doi: 10.1099/ijs.0.02307-0

Sievers, F., & Higgins, D.G. (2018) Clustal Omega for making accurate alignments of many protein sequences. Protein Science, 27, 135–145. doi: 10.1002/pro.3290

Smith, E.A., & Macfarlane, G.T. (1996). Enumeration of human colonic bacteria producing phenolic and indolic compounds: effects of pH, carbohydrate availability and retention time on dissimilatory aromatic amino acid metabolism. Journal of Applied Bacteriology, 81, 288–302. doi: 10.1111/j.1365-2672.1996.tb04331.x

Trott, O., & Olson, A.J. (2010). AutoDock Vina: improving the speed and accuracy of docking with a new scoring function, efficient optimization, and multithreading. Journal of Computational Chemistry, 31, 455–461. doi: 10.1002/jcc.21334

Venskutonytė, R., Koh, A., Stenström, O., Khan, M.T., Lundqvist, A., Akke, M., … Lindkvist-Petersson, K. (2021). Structural characterization of the microbial enzyme urocanate reductase mediating imidazole propionate production. Nature Communications, 12, 1347. doi: 10.1038/s41467-021-21548-y.

Wargnies, M., Plazolles, N., Schenk, R., Villafraz, O., Dupuy, J.W., Biran, M., … Bringaud, F. (2021). Metabolic selection of a homologous recombination-mediated gene loss protects Trypanosoma brucei from ROS production by glycosomal fumarate reductase. Journal of Biological Chemistry, 296, 100548. doi: 10.1016/j.jbc.2021.100548

