## Supplemental figures 1-3 for "A redox-regulated, heterodimeric NADH:cinnamate reductase in *Vibrio ruber*"

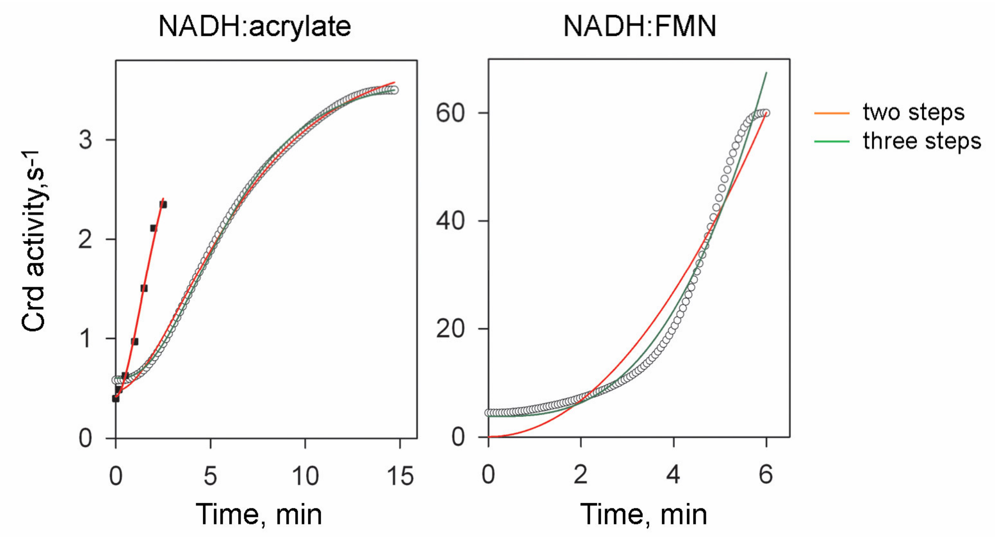


**Figure S1.** The time dependencies of Crd activity derived from the time-courses of anaerobic acrylate and FMN reduction. Small circles show the activity values determined at 100 time points as the slope of the tangent (-*d*[NADH]/*d*t) to the blue curves on Fig. 6C and 6D (reactions started by Crd addition) using MATLAB (The MathWorks, Inc.), as for Fig. 6B. Squares refer to the activity values directly estimated upon acrylate addition to Crd preincubated for the indicated time with NADH. A home-made low dead-time cuvette was used for kinetic measurements in this case; other conditions were as for Fig. 6C. The red and green lines show the best fits for the kinetic models involving two and three consecutive activation steps, respectively.

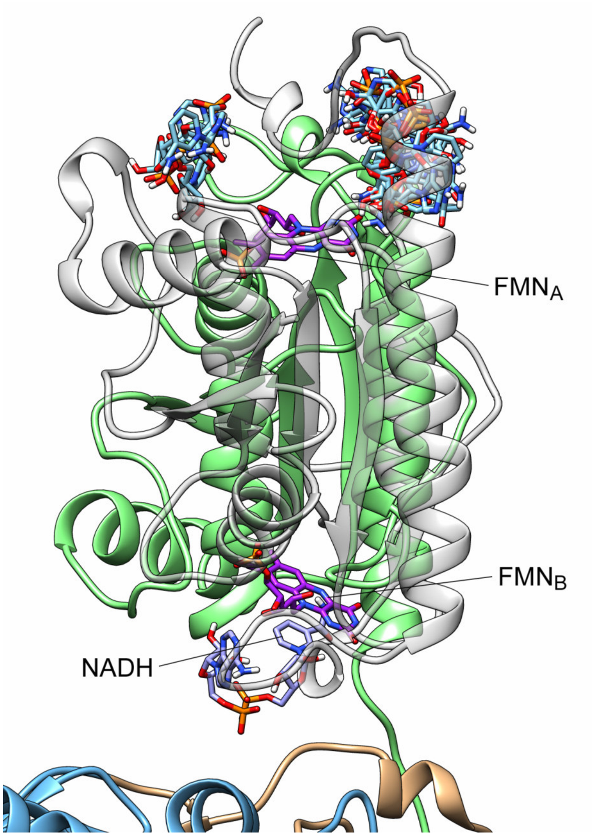


**Figure S2.** NADH docking to the Crd d1 fragment in the vicinity of the bound flavin groups. A single position in which NADH molecule stacks FMN$_B$ is shown. No such position was found for FMN$_A$; instead, nine best-score positions formed two non-stacking ensembles. Domain colors are as in Fig. 8.

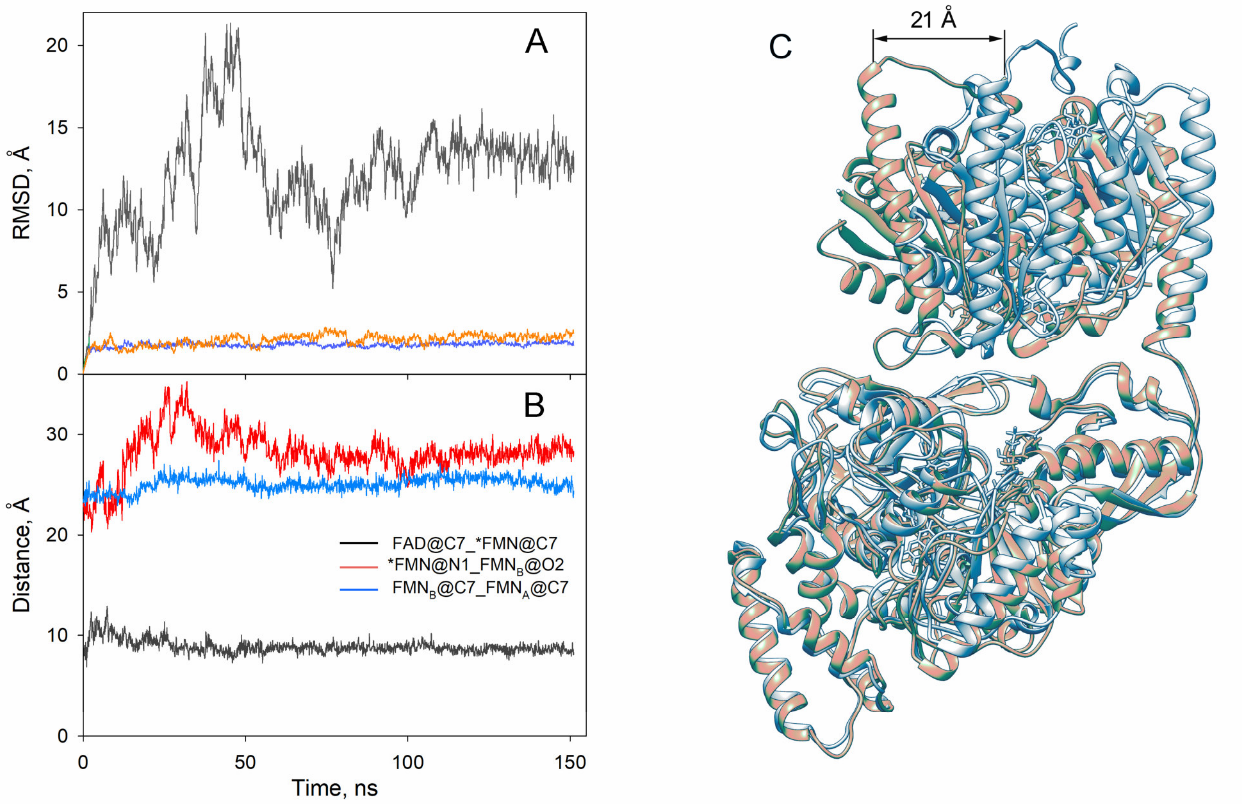


**Figure S3.** The refinement of the flavin-bound AlphaFold-predicted Crd structure by molecular dynamics simulations in explicit solvent. The system was hydrogenated, electroneutralized by adding 170 $K^+$ and 125 $Cl^-$ ions (corresponding to 100 mM KCl concentration) using the program TLEAP of the AMBER 22 package (http://ambermd.org/) and solvated with a 12 Å OPC water molecules (the "first shell of water molecules" was made using 3D-RISM and METATWIST programs of the Amber 22 package) in the final box of approximately 2,200,000 $Å^3$ volume. The ff19SB force field was used for protein residues (Tian et al., 2019; http://ambermd.org). The general AMBER force field was used for the FAD and FMN residues and the FMN-binding serine residue (Ser257). Partial charges for ligands were calculated using ANTECHAMBER program with AM1-bcc options. The total number of atoms in the simulations was approximately 200,000. The system was minimized using a SANDER function with 3,000 steps of steepest descent and 2,000 steps of conjugate gradient with the restraint of 10 kcal·mol$^{-1}$·Å$^{-2}$ on the protein and ligands atoms. The system was gradually heated to 300 K under constant volume conditions with 10 kcal·mol$^{-1}$·Å$^{-2}$ restraints on the protein and ligand atoms for 25,000 1-fs steps, then restraints were gradually removed in seven stages with 50,000-100,000 1–2-fs steps. The MD simulations were performed using the PMEMD program (CUDA implementation) of the AMBER 22 suite under constant pressure (NPT ensemble) at 300 K using a Langevin thermostat and piston, for temperature and pressure control, respectively. Long-range electrostatic interactions were computed using Particle Mesh Ewald (PME) algorithm. The 10-Å cut-off production MD was performed for 150 ns, with snapshots saved every 10 ps. Post-processing trajectory analyses were carried out with the program CPPTRAJ of the AMBER 22 suite.

(A) Changes in the backbone-atom RMSD during the MD simulations: blue, for the *NADH:flavin* domains (d1 parts) only; red, for the *FAD binding 2* and *FMN bind* domains only; black, for the *NADH:flavin* domains (d1 parts) only in the structures superimposed by their *FAD binding 2* and *FMN bind* domains, as in panel C. (B) Changes in the distances between the flavin groups during the MD simulations. The distances are between the atoms indicated in the panel label. (C) The AlphaFold2-generated Crd structures before and after MD simulations (blue and orange, respectively) superimposed by their *FAD binding 2* and *FMN bind* domains.

**Reference**

Tian, C., Kasavajhala, K., Belfon, K.A.A., Raguette, L., Huang, H., Migues, A.N., Bickel, J., Wang, Y., Pincay, J., Wu, Q., & Simmerling, C. (2019). ff19SB: Amino-acid-specific protein backbone parameters trained against quantum mechanics energy surfaces in solution. J. Chem. Theory Comput. **16**, 528–552.
